# The histone demethylase JMJ27 acts during the UV-induced modulation of H3K9me2 landscape and facilitates photodamage repair

**DOI:** 10.1101/2024.04.01.587525

**Authors:** Philippe Johann to Berens, Jackson Peter, Sandrine Koechler, Mathieu Bruggeman, Sébastien Staerck, Jean Molinier

## Abstract

Plants have evolved sophisticated DNA repair mechanisms to cope with the deleterious effects of UV-induced DNA damage. Indeed, DNA repair pathways cooperate with epigenetic-related processes to efficiently maintain genome integrity. However, it remains to be deciphered how photodamages are recognized within different chromatin landscapes, especially in compacted genomic regions such as constitutive heterochromatin. We combined cytogenetics and epigenomics to identify that UV-C irradiation induces modulation of the main epigenetic mark found in constitutive heterochromatin, H3K9me2. We demonstrated that the histone demethylase, Jumonji27 (JMJ27), is responsible for the UV-induced reduction of H3K9me2 content at chromocenters. In addition, we identified that JMJ27 forms a complex with the photodamage recognition factor, DNA Damage Binding protein 2 (DDB2), and that the fine tuning of H3K9me2 contents orchestrates DDB2 dynamics on chromatin in response to UV-C exposure. Hence, this study uncovers the existence of an interplay between photodamage repair and H3K9me2 homeostasis.

## INTRODUCTION

Sunlight is used by plants as a source of energy and as a signal to trigger developmental transitions. Nevertheless, plants have to cope with ultraviolet radiations (UV) that damage DNA. The genotoxic effect of UV leads to the formation of Cyclobutane Pyrimidine Dimers (CPDs) and 6–4-Photoproducts (6–4 PPs) occurring between di-pyrimidines (Britt, 1995). In plants, several photodamage repair pathways are mobilized to either photoreactivate or excise UV-induced DNA lesions (Molinier, 2017). CPDs and 6-4 PPs are preferentially repaired by a light-dependent error-free mechanism, called Direct Repair (DR) involving photolyases (Britt, 1995). Two active photolyases are found in *Arabidopsis thaliana*, PHOTOLYASE 1 (PHR1) and UV RESISTANCE 3 (UVR3) photoreactivating CPDs and 6-4 PPs, respectively (Britt, 1995).

A light-independent repair process, called Nucleotide Excision Repair (NER), also exists (Molinier, 2017). NER contains two sub-pathways, the Transcription-Coupled Repair (TCR) and the Global Genome Repair (GGR; Schärer, 2013), that enzymatically remove photodamage along actively transcribed DNA strands or throughout the genome, respectively (Schärer, 2013). In transcribed genomic regions, photolesions block transcription and the stalled RNA polymerase II triggers the recognition signal to initiate TCR (Lainé and Egly, 2006) whilst the DNA DAMAGE-BINDING protein 2 (DDB2) recognizes photodamage for GGR in poorly transcribed/untranscribed regions (Chu and Chang, 1988).

Thus, the permissive or the suppressive epigenetic landscapes of the photodamaged genomic regions define by which NER sub-pathway the lesions will be detected (Johann to Berens and Molinier, 2020). The mechanisms by which GGR recognizes photodamaged DNA sequences wrapped around nucleosomes are still under investigation. Cryo-Electron Microscopy approaches demonstrated how human DDB2 senses photolesions on nucleosomes (Matsumoto *et al*., 2019). The human DDB2-containing complex allows the recruitment of chromatin remodelers (Zhao *et al*., 2009; Jiang *et al*., 2010; Pines *et al*., 2012), of histone writer (Balbo Pogliano *et al*., 2017; Zhu *et al*., 2018) and displaces histone H1 (Fortuny *et al*., 2021). In Drosophila, NER is enhanced by H3K9me3 demethylation of heterochromatin involving a Lysine-9 demethylase (Palomera-Sanchez *et al*., 2010) that most likely triggers remodeling/relaxation (Shu et al., 2012).

Whereas in human and drosophila, constitutive heterochromatin is enriched in H3K9me3 (Burton *et al*., 2020; Wei *et al*., 2021), in Arabidopsis, H3K9m2 is predominant (Xu and Jiang, 2020). H3K9me2 is deposited by SUVH4 (Suppressor of Variegation 3-9 Homolog Protein 4 also called KRYPTONITE: KYP) and to some extent by SUVH5 and 6 (Jackson *et al*., 2002; Ebbs and Bender, 2006; Li *et al*., 2018). H3K9me2 transcriptionally silences transposable elements (TE; Law and Jacobsen, 2010) and regulates expression of several protein coding genes (PCG; Saze *et al*., 2008; Miura *et al*., 2009). Arabidopsis carries also 3 H3K9me2 demethylases with Jumonji (JMJ) C domain-containing: JMJ29, JMJ25 and JMJ27 (Fan *et al*., 2012; Dutta *et al*., 2017; Hung *et al*., 2020). Whereas JMJ29 was described as a regulator of trichome development, JMJ25, also called INCREASE IN BONSAI METHYLATION 1 (IBM1) prevents heterochromatinization of genic regions (Saze *et al*., 2008; Miura *et al*., 2009), regulates plant immunity (Chan and Zimmerli 2019) and RNA-directed DNA methylation (RdDM) pathway (Fan *et al*., 2012). JMJ27 coordinates defense response against pathogens, controls flowering time (Dutta *et al*., 2017), drought stress response (Wang *et al*., 2021a) and acts during meiosis together with IBM1 (Cheng *et al*., 2022).

Although the roles of histone H3K9me2 methyltransferases/demethylases in the control of transposition and gene expression have been characterized (Saze *et al*., 2008; Law and Jacobsen, 2010), the putative interconnections between H3K9me2 homeostasis and maintenance of genome integrity (*i.e.,* DNA repair) remain to be elucidated.

In this study we combined different approaches to uncover how photodamage repair and H3K9me2 homeostasis are interconnected. Using cytogenetics and deep-learning-based tools, we found that photolesions are present at chromocenters and that UV-C irradiation induces the modulation of H3K9me2 contents in these regions. We identified that H3K9me2 dynamics relies on both DDB2 and JMJ27. Chromatin immunoprecipiation (ChIP) experiments allowed identifying genomic regions exhibiting loss of H3K9me2 content upon UV-C irradiation in a JMJ27-dependent manner. Finally, we found that KYP and JMJ27 act in the GGR pathway and influence the UV-C-induced spatio-temporal loading and release of DDB2 on chromatin. Altogether, our results document the existence of an interplay between H3K9me2 homeostasis and photodamage repair.

## RESULTS

### UV-C irradiation induces modulation of H3K9me2 at chromocenters

In order to determine the sub-nuclear distribution of UV-C-induced photoproducts we used anti-CPDs and anti-6-4-PPs antibodies to immunolabel WT nuclei. Upon irradiation, both CPDs and 6-4 PPs fluorescent signals are found to be located in the nucleoplasm and to overlap with DAPI labeled chromocenter regions (Fig. 1). Given that photodamage repair factors have to access photolesions within the compacted chromocenters (Dabin *et al*., 2023) we checked whether UV-C irradiation triggers modulation of constitutive heterochromatin histone marks: H3K9me2 and H3K27me1. We specifically trained the deep-learning-based tool, Nucl.Eye.D (Johann to Berens *et al*., 2022) for signal recognition of H3K9me2 and H3K27me1. This allowed the automated segmentation of chromocenter-like structures exhibiting H3K9me2 and H3K27me1 signals and their accurate quantifications.

**Figure 1:**
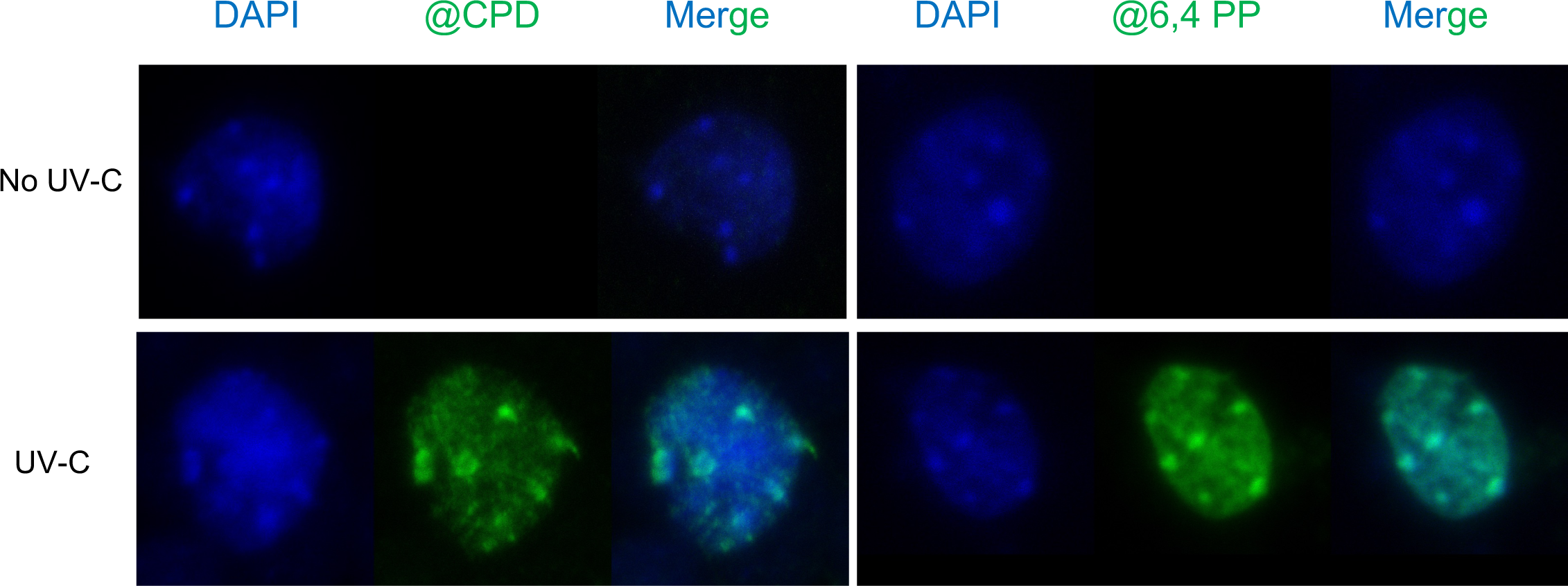
Nuclear of localization of UV-C-induced photodamage. Immunolabeling of CPD and 6,4-PP (green) on DAPI stained (Blue) WT (Col-0) nuclei prepared prior (no UV-C) and 30 min upon UV-C exposure. Scale bar = 5μm.

We determined H3K9me2 and H3K27me1 occupancies prior and upon UV-C exposure. In WT plants, both H3K9me2 and H3K27me1 signals are, as expected, located at chromocenters (Fig. 2A). These signals are, reduced, even absent, in both *kyp suvh-5,6* (H3K9me2 methyltransferases) and *atxr-5,6* (H3K27me1 methyltransferases) mutant plants (Fig. S1A). Two hours upon UV-C exposure, the H3K9me2 content significantly decreases in WT plants (Fig. 2B). These differences are mainly due to changes in occupancy, reflecting reshaping, whereas H3K9me2 relative intensity remains stable (Fig. S1B, C). 24h upon irradiation the H3K9me2 content is re-established and is even higher than prior irradiation (Fig. 2B). Conversely, H3K27me1 contents do not exhibit significant changes upon UV-C irradiation (Fig. 2C). We can observe at 24h that the H3K27me1 occupancy decreases and the relative intensity increases whereas comparable content values are detected (Fig. S1D, E). Altogether, these data demonstrate that UV-C alters only H3K9me2 contents, and that these changes are likely due to chromocenters reshaping.

**Figure 2:**
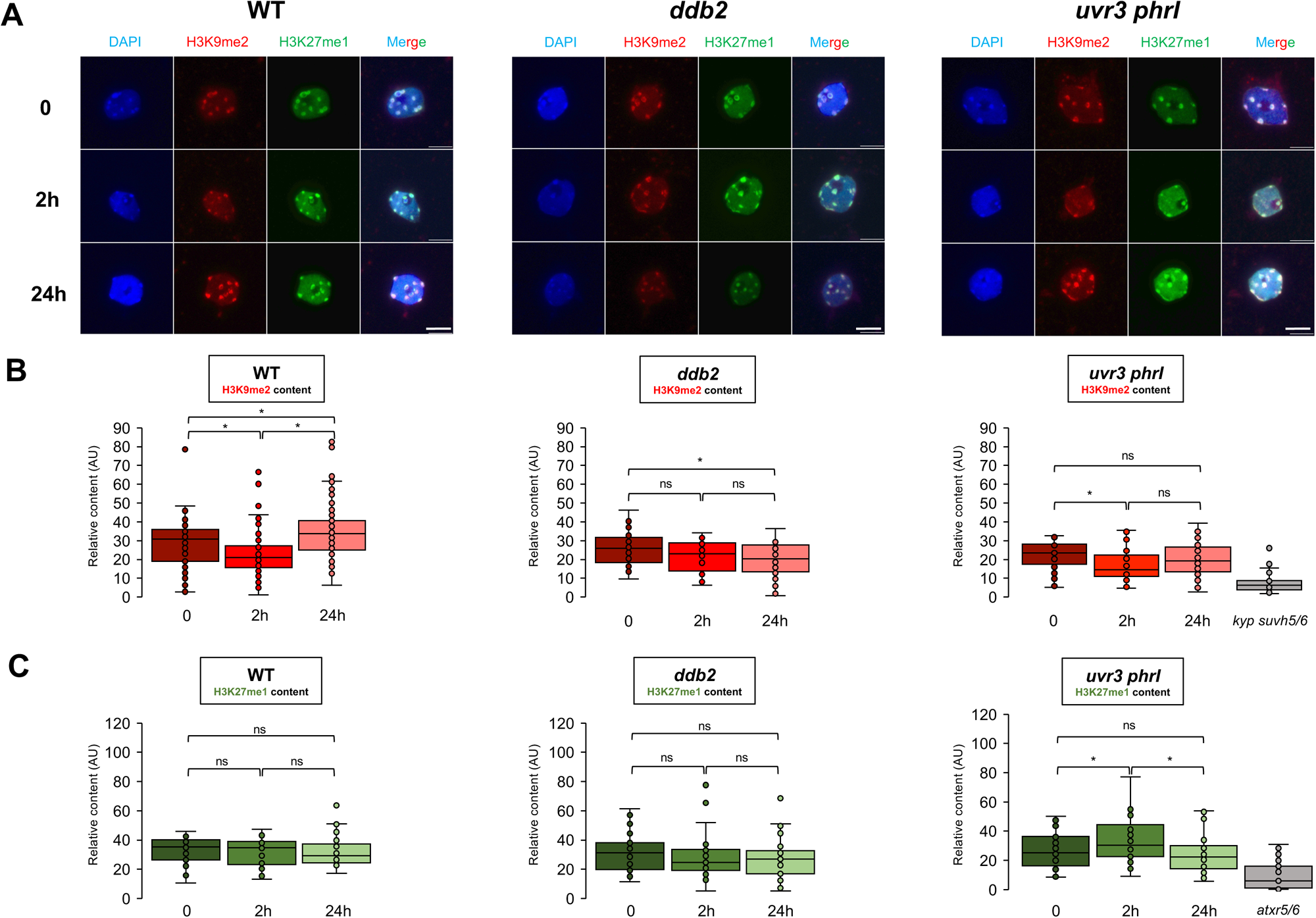
H3K9me2 and H3K27me1 patterns in WT and photodamage deficient plants upon UV-C exposure. **A.** Microscopy images of DAPI, H3K9me2 and H3K27me1 immuno-stained arabidopsis nuclei isolated from WT (Col-0), *ddb2* and *uvr3phr1* leaves in control condition (0) and upon UV-C irradiation (2h, 24h). Scale bar = 5μm. **B.** Box plots of the H3K9me2 Content (Occupancies * Relative Intensities) in a population of at least 30 nuclei per condition. Each dot represents the measure for 1 nucleus. *kypsuvh5/6* plants were used as control. Mann Whitney Wilcoxon test \**p* < 0.01; \*\**p*< 0.05 ns: nonsignificant. **C.** Same as **B.** for and H3K27me. *atxr5/6* plants were used as control.

To further investigate if these changes are related to photodamage repair pathways, immunostaining of H3K9me2 and H3K27me1 marks have been performed on DAPI stained nuclei prepared from plants deficient for GGR (*ddb2*) and DR (*uvr3phr1*). In *ddb2* plants, in which the DR pathway would be mainly used to repair photoproducts, the H3K9me2 content remains stable at 2h (Fig. 2B) and exhibits a significant lower content only 24h upon UV-C (Fig. 2B). H3K27me1 content remains stable like in WT plants (Fig. 2C). This suggests that the UV-C-induced modulation of H3K9me2 content depends on DDB2 and that alternative DNA repair pathways (*i.e.*, DR) do not even trigger such change.

In *uvr3phr1* plants, GGR is thought to be predominantly used to repair photodamage. The kinetics of H3K9me2 content is closely related to the one observed in WT plants (Fig. 2B). At 24h, the H3K9me2 content reaches again the level observed in control condition (0) but does not show further increase as observed in WT plants (Fig. 2B). H3K27me1 contents transiently increase, suggesting that GGR likely triggers deposition and/or that DR prevent modulation of this epigenetic mark (Fig. 2C).

These analyses highlight that UV-C irradiation alters only H3K9me2 pattern at chromocenters and that DDB2 is involved in this dynamic. Hence, GGR might interplay with factors modulating H3K9me2 landscape.

### JMJ27 contributes to the UV-C-induced modulation of H3K9me2 content at chromocenters

H3K9me2 dynamics within chromocenters can rely on different processes such as chromatin remodeling (Zhao *et al*., 2009; Jiang *et al*., 2010; Pines *et al*., 2012), histone displacement (Fortuny *et al*., 2021), erasure-writing of post-translational modification (PTM) (Palomera-Sanchez *et al*., 2010; Balbo Pogliano *et al*., 2017; Zhu *et al*., 2018). We tested the putative role of active histone demethylation in the alteration of H3K9me2 contents at chromocenters upon UV-C exposure. We used Nucl.Eye.D to quantify the H3K9me2 signal in nuclei prepared from *ibm1* and *jmj27* plants defective for the expression of the main Arabidopsis H3K9me2 demethylases (Saze *et al*., 2008; Dutta *et al*., 2017). Both WT and *ibm1* plants exhibit a significant decrease of H3K9me2 occupancy/content at 2h whilst the H3K9me2 signal remains stable in *jmj27* plants (Fig. 3). The lack of H3K9me2 decay at chromocenters in UV-C-treated *jmj27* plants suggests that JMJ27 is involved in the removal of this histone PTM.

**Figure 3:**
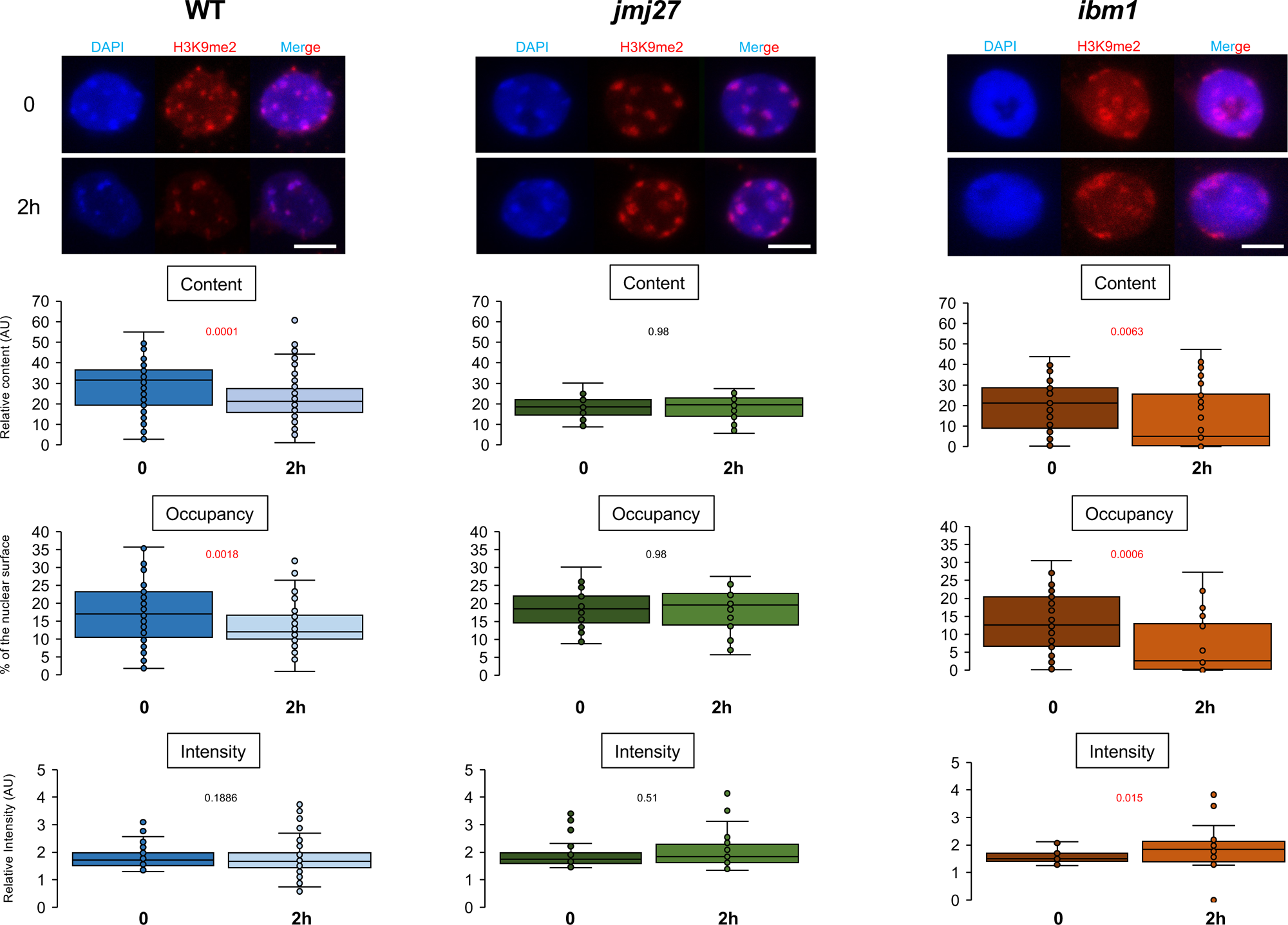
Measurements of H3K9me2 signals in WT, *jmj27* and *ibm1* mutant plants upon UV-C exposure. **A.** Microscopy images of DAPI and H3K9me2 immuno-stained arabidopsis nuclei isolated from WT*, jmj27* and *ibm1* leaves in control condition (0) and upon UV-C irradiation (2h). Scale bar = 5μm. **B.** Box plots of H3K9me2 Content (Occupancies * Relative Intensities). **C.** Box plots of H3K9me2 Occupancy (percent of the nuclear surface occupied by chromocenter like structure). **D.** Box plots of H3K9me2 Relative Intensity (ratio of mean chromocenter structure intensity / Mean nuclei intensity). A population of at least 25 nuclei per condition was used. Each dot represents the measure for 1 nucleus. Exact p values are shown (Mann Whitney Wilcoxon test).

### UV-C-induced alterations of H3K9me2 landscape

In order to gain in resolution and to identify genomic regions exhibiting modulation of H3K9me2 contents in a UV-C- and/or JMJ27-dependent manner, we performed H3K9me2 ChIP using WT- and *jmj27-*irradiated plants.

Prior irradiation, WT plants exhibit an enrichment of H3K9me2 at more than 1000 genomic regions, predominantly located at centromeres and pericentromeres (Fig. 4A, S2). Upon irradiation the number of H3K9me2-enriched regions decreases in WT plants whereas in *jmj27* plants, it transiently increases (Fig. 4A). Interestingly, the number of H3K9me2-containing regions in *jmj27*-untreated plants is lower than in WT plants (Fig. 4A). We found that *jmj27* plants accumulated higher levels of both short and long *IBM1* mRNA isoforms (Rigal *et al*., 2012; Saze *et al*., 2013; Fig. S3A). This holds true upon UV-C exposure (Fig. S3B). Thus, it is likely that regulatory interplays between JMJ27 and IBM1 exist to fine tune the H3K9me2 landscape.

**Figure 4:**
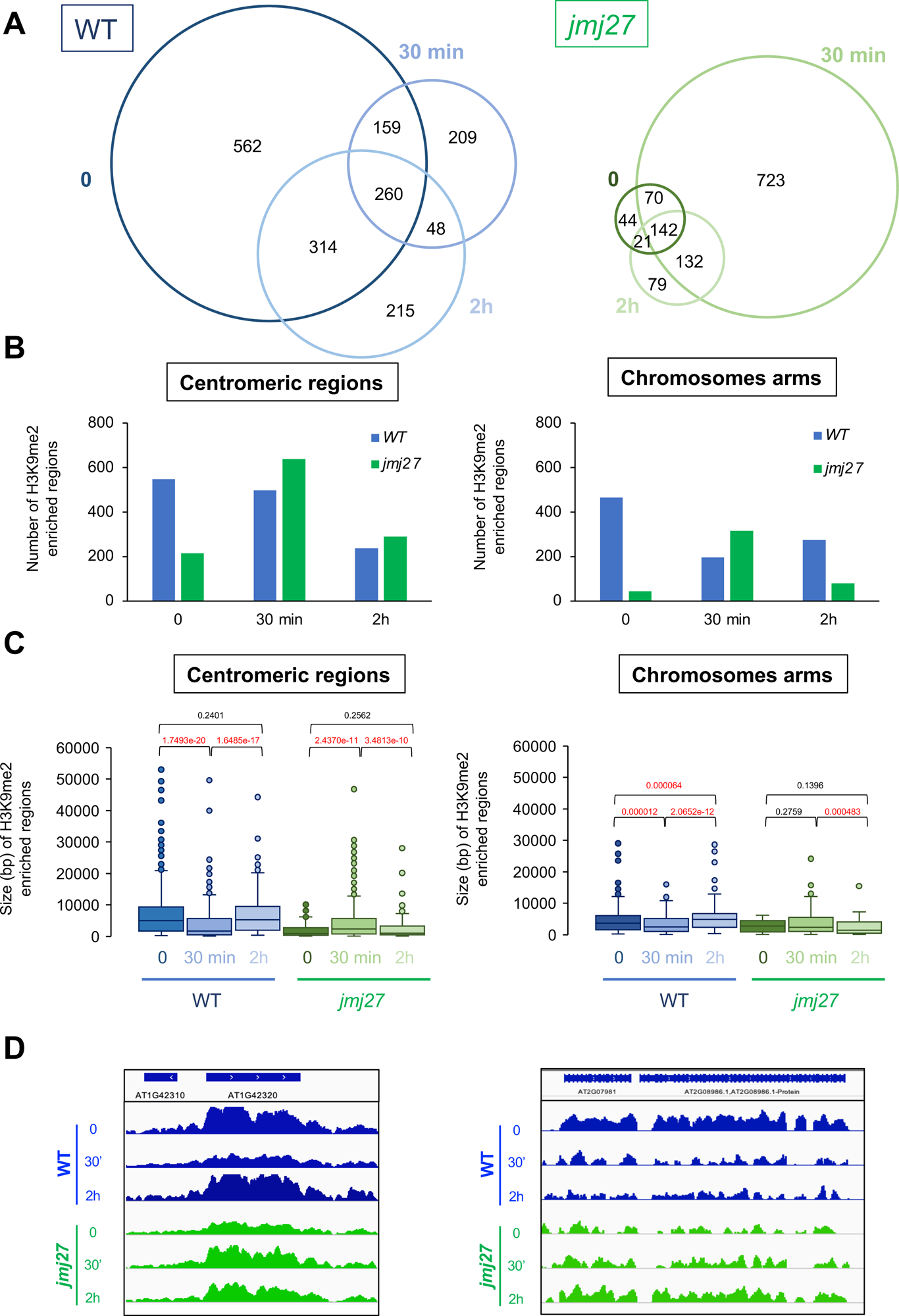
H3K9me2-enriched genomic regions in WT and *jmj27* mutant plants upon UV-C exposure. **A.** Venn diagram showing the overlap between H3K9me2-enriched genomic regions in WT (left) and *jmj27* mutant plants (right) prior (0), 30 min and 2h upon UV-C exposure. **B.** Histogram showing the number of significantly H3K9me2-enriched regions in centromeric regions (left panel) and in chromosomes arms (right panel) prior UV-C exposure (0), 30 min and 2h upon irradiation in WT (blue) and *jmj27* (green) plants. **C.** Box plots representing the size (bp) of H3K9me2-enriched regions in centromeric regions (left panel) and in chromosomes arms (right panel) prior UV-C exposure (0), 30 min and 2h upon irradiation in WT (blue) and *jmj27* (green) plants. Exact p values are shown (Mann Whitney Wilcoxon test). **D.** Genome browser tracks showing examples of H3K9me2 contents in WT (blue) and *jmj27* (green) plants.

The proportions of PCG, TE and intergenic regions containing H3K9me2 do not significantly vary between WT and *jmj27* plants along the time course (Fig. S4A, B) highlighting that UV-C exposure does not lead to a preferential redistribution of H3K9me2 at particular genetic entities. Interestingly, the LTR/Gypsy and DNA superfamilies represent a large proportion of TE exhibiting H3K9me2 enrichment in both WT and *jmj27* plants (Fig. S4C, D). In order to better characterize the UV-C-induced changes of H3K9me2 landscape, we uncoupled the analysis between centromeres and chromosomes arms. In WT plants the number of H3K9me2 regions overlapping with centromeres decreases at 2h (Fig. 4B). Their sizes transiently drop and reach back at 2h (Fig. 4C). Conversely, in *jmj27* plants, the number and the size of the H3K9me2-enriched regions transiently increase (Fig. 4B-D). This shows that UV-C induces H3K9me2 dynamics at chromocenters and that JMJ27 is likely involved in this process. In chromosomes arms, the number and of the sizes of H3K9me2-containing regions decrease at 30 min in WT plants and are suppressed in *jmj27* plants (Fig. 4B, C, D).

Altogether these results show that the kinetics of H3K9me2 dynamics differs between centromeres and chromosomes arms. Moreover, the transient changes of H3K9me2 contents underpin the existence of UV-C-induced loss and *de novo* deposition of H3K9me2.

The comparative analysis (WT *vs jmj27*) of our ChIP data allowed identifying genomic regions displaying an UV-C-induced and a JMJ27-dependent loss of H3K9me2. These regions overlap mainly with TE (Fig. S5A) whose LTR/Gypsy and DNA superfamilies represent a significant proportion (Fig. S5B). Thus, our results show that JMJ27 preferentially targets TEs in response to UV-C irradiation conversely to IBM1 that predominantly targets PCG (Saze *et al*., 2008). Finally, these analyses show that UV-C irradiation triggers modulation of H3K9me2 landscape genome wide and reinforced the idea that JMJ27 plays a role in this reshaping.

### KYP and JMJ27 mutant plants are defective in 6-4 PPs repair

Given that UV-C irradiation modulates H3K9me2 contents at chromocenters where photolesions are formed, it might be assumed that H3K9me2 homeostasis could contribute to their repair. To test this, we measured repair efficiency of JMJ27 and KYP defective plants, using dot blot (Schalk *et al*., 2017). In WT plants, around 60% of the UV-C-induced CPDs remain detected whereas only 20% and 5% are still present in *jmj27* and *kyp* plants, respectively (Fig. 5A). This shows that CPD repair is not impaired in JMJ27 and KYP deficient plants. Conversely, higher amounts of 6-4 PPs are detected in both *jmj27* and *kyp* plants compared to WT plants (Fig. 5B). These results show that JMJ27 and KYP are important for efficient repair of one type of photolesions: 6-4 PPs, and highlight that H3K9me2 homeostasis and photodamage repair are interconnected.

**Figure 5:**
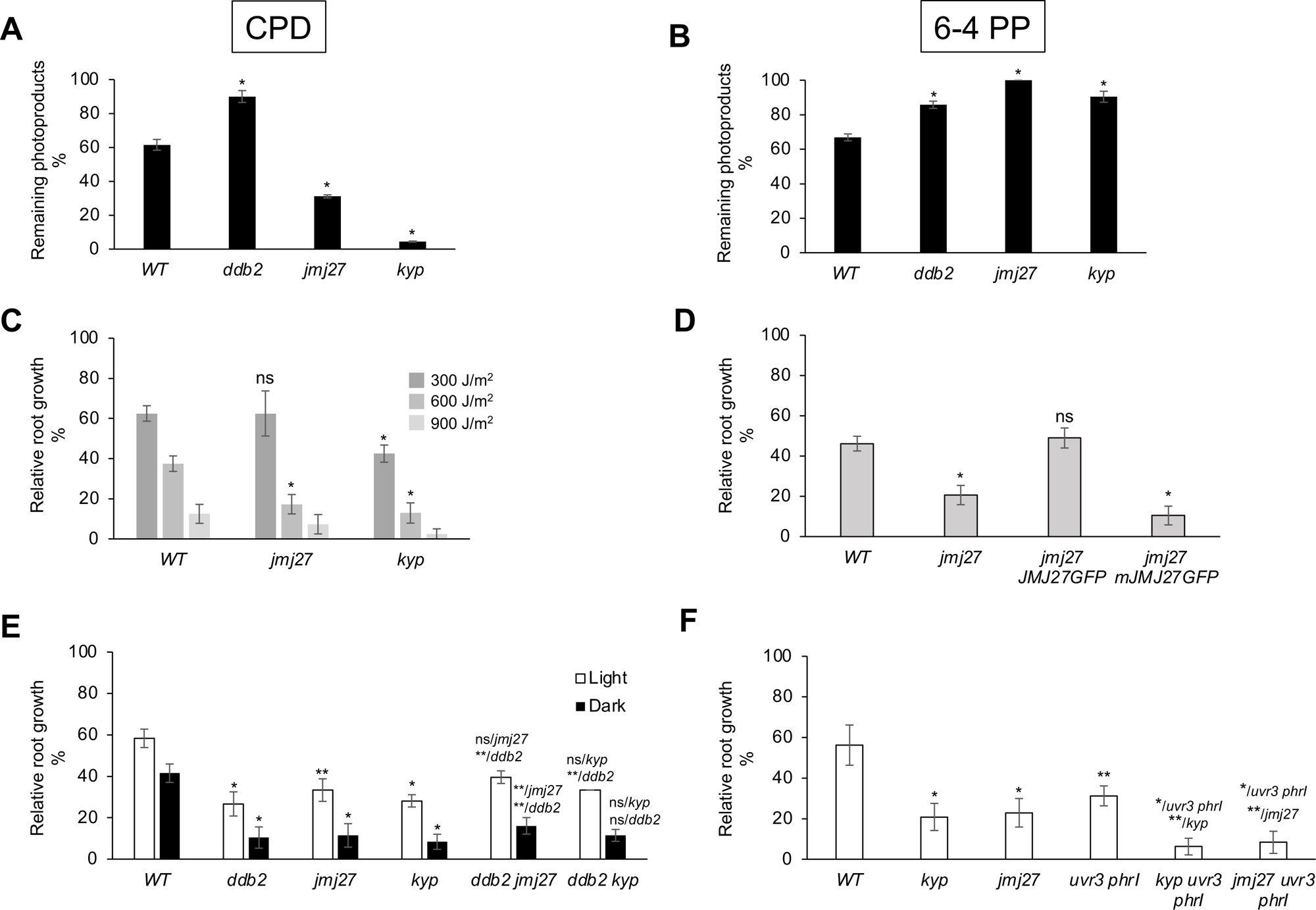
Photodamage repair assay and genetic interactions of *JMJ27*, *KYP* and photolesions repair mutations. Photodamage repair assay. Histogram representing the relative amounts of **(A)** CPDs and **(B)** 6-4 PPs 1h upon UV-C irradiation (± SD) in WT, *ddb2, jmj27* and *kyp* plants. *ddb2* is a photodamage deficient mutant and was used as positive control. Photodamage repair impairment is displayed by a higher relative amount of CPD and/or 6-4 PP than WT plants 1h upon UV-C exposure. Intensity of each dot was quantified and normalized to that of CPDs and 6-4 PPs at time 0 to calculate the remaining photodamage content after 1 h. Twenty plants per replicate were used, and experiments were duplicated. *t* test \**p* < 0.01. **C.** Root growth assay. Seven-day-old WT, *JMJ27* and *KYP* mutant plants were exposed to increasing doses of UV-C. Root growth was calculated relative to the corresponding untreated plants (±SD). **D.** Complementation assay. Seven-day-old WT, *JMJ27* mutant plants and *jmj27* plants expressing either the JMJ27 histone methyltransferase fusion protein (JMJ27-GFP) or the catalytic defective JMJ27 histone methyltransferase fusion protein (mJMJ27-GFP) were exposed to UV-C. *t* test \**p* < 0.01; ns: nonsignificant. **E.** Genetic interactions between *ddb2*, *kyp* and *jmj27* grown under light or dark recovery conditions upon UV-C exposure. *t* test \**p* < 0.01; \*\**p* < 0.05; ns: nonsignificant compared with the corresponding WT plants or the single mutant plant. **F.** Same as **E.** for *DR (UVR3PHRI)*, *KYP* and *JMJ27* defective plants. Seven-day-old WT, *kyp, jmj27*, *uvr3phrI* plants and triple mutant plants (*kyp*-*uvr3phrI*, *jmj27*-*uvr3phrI*) were exposed to UV-C. *t* test \**p* < 0.01**; *p* < 0.05 compared with the corresponding single/double mutants for triple mutants. Eight plants per replicate were used, and three independent biological replicates were performed.

### KYP and JMJ27 act in GGR

In order to better define how H3K9me2 homeostasis and DNA repair interconnect, we analyzed the genetic interactions between *KYP*, *JMJ27* and photodamage repair mutations. Both *jmj27* and *kyp* mutant plants exhibit UV-C hypersensitivity compared to WT plants (Fig. 5C). In *jmj27 JMJ27-GFP* line this phenotype is suppressed, demonstrating the functional complementation (Fig. 5D, S6A). Moreover, immunoblot analysis of H3K9me2 content of irradiated *jmj27 JMJ27-GFP* line shows, a decrease of this PTM at 2h (Fig. S6B). Conversely, *jmj27* plants expressing a catalytic deficient JMJ27 protein (mJMJ27-GFP; Wang *et al*., 2021b), exhibit UV-C hypersensitivity and accumulation of H3K9me2 at 2h (Fig. 5D, S6A, B). These results suggest that JMJ27 and active H3K9me2 demethylation, are important for photodamage repair. In order to determine in which repair pathway JMJ27 and KYP act, we studied the genetic interactions between *jmj27*, *kyp,* GGR (*ddb2*) and DR (*uvr3phr1*) mutations. When *ddb2* is combined with *jmj27* or *kyp*, the UV-C sensitivity does not differ from the one observed in single mutant plants (Fig. 5E) suggesting that *ddb2*, *jmj27* and *kyp* mutations are epistatic. Thus, KYP and JMJ27 act in the GGR pathway. *kyp* and *jmj27* double mutant plants exhibit an epistatic interaction, showing that they act in the same pathway (Fig. S7). In order to determine whether KYP and JMJ27 are also involved in DR, we produced *uvr3phr1 kyp* and *uvr3phr1 jmj27* triple mutant plants. Both combinations show synergistic effects (Fig. 5F) ruling out that KYP and JMJ27 act in the DR. Under dark grown recovery condition, preventing photodamage reversion by photolyases, similar synergistic effects are observed (Fig. 5E). These data strengthen the findings that KYP and JMJ27 are predominantly involved in GGR.

### H3K9me2 content influences UV-induced internucleosomal-nucleosomal DDB2 dynamics

We identified that UV-C irradiation triggers a rapid decrease of H3K9me2 content in a DDB2- and JMJ27-dependent manner, suggesting that both factors might act concomitantly. Therefore, we tested the kinetics of JMJ27 and DDB2 loading on chromatin upon UV exposure. For this we conducted immunoblot analyses of chromatin-enriched fraction (insoluble) of *jmj27* plants expressing either JMJ27-GFP (complemented line) or mJMJ27-GFP (catalytic deficient demethylase; Wang *et al*., 2021a). Both JMJ27-GFP and DDB2 load on chromatin at 30 min upon UV-C exposure (Fig. 6A). Conversely, the catalytic deficient mJMJ27-GFP fusion protein remains enriched in chromatin fraction during the time course whereas DDB2 displays a progressive release (Fig. 6A).

**Figure 6:**
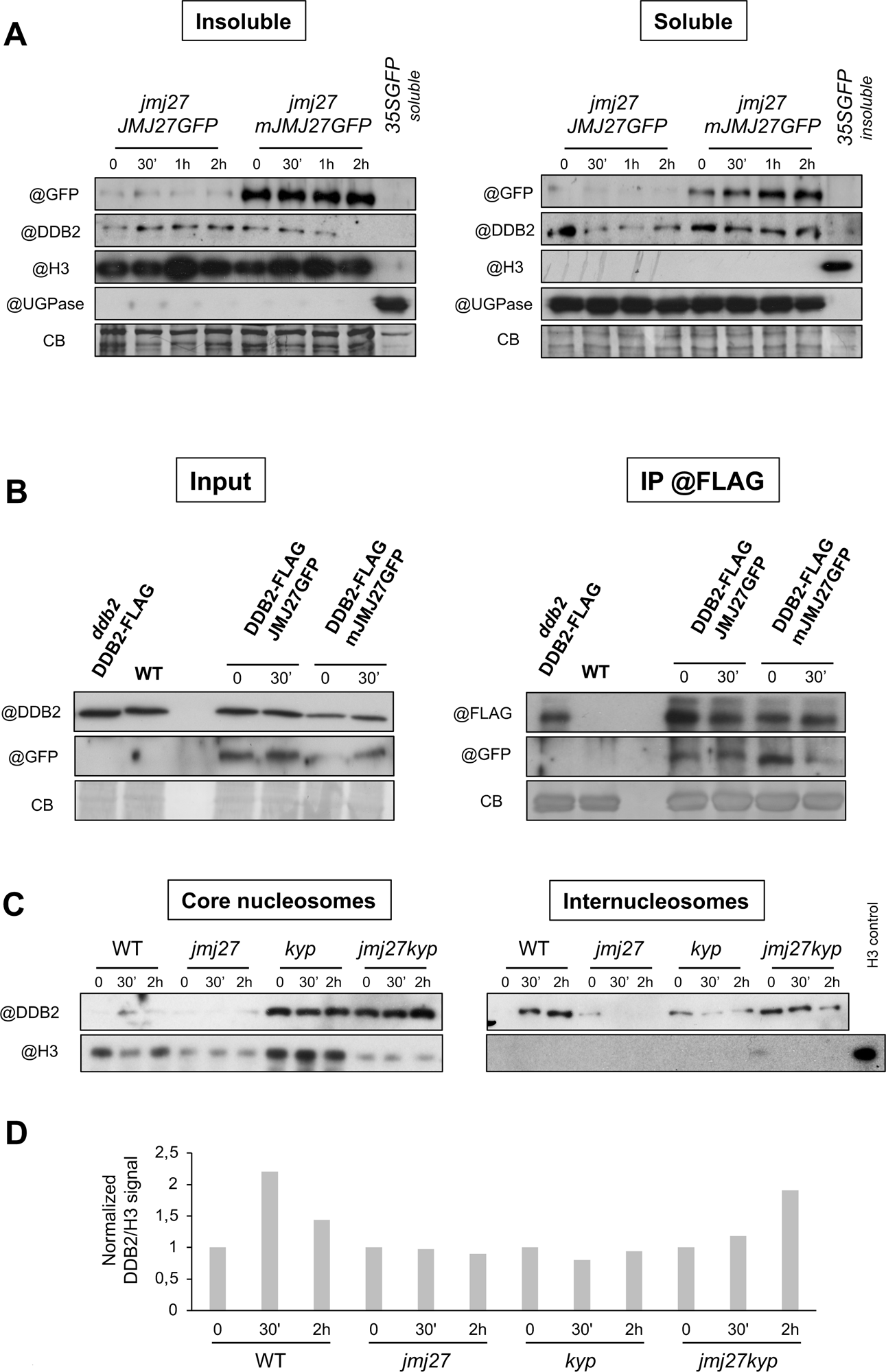
DDB2 contents at internucleosomal and nucleosomal sites and DDB2-JMJ27 complex. **A.** Immunoblot analysis of JMJ27-GFP and DDB2 contents in insoluble (chromatin-enriched) and soluble fractions prepared from *jmj27 JMJ-GFP* and *jmj27 mJMJ-GFP* plants prior (0), 30 min, 1h and 2h upon UV-C exposure. The anti-H3 antibody was used as positive control for insoluble fraction and the anti-UGPase antibody was used as positive control for soluble fraction. 35S-GFP plants were used as negative control for anti-GFP signal at 125 kDa corresponding to both JMJ27-GFP and mJMJ27-GFP fusion protein. Coomassie blue staining (CB) of the blot is shown for each fraction. **B.** *In vivo* pull-down of JMJ27-GFP with DDB2-FLAG protein prior (0) and 30 min upon UV-C exposure. *ddb2-2*/*DDB2*-*FLAG* expressing plants crossed with either JMJ27-GFP or mJMJ27-GFP (mutated catalytic methyltransferase site) were used for IP assays using anti-FLAG antibody. DDB2-FLAG and WT plants were used as negative control for GFP signal. Coomassie blue staining (CB) of the blot is shown. **C.** Immunoblot analysis of DDB2 content at nucleosomal and internucleosomal sites in WT, j*mj27*, *kyp* and *jmj27kyp* plants prior (0), 30 min and 2h upon UV-C exposure. Anti-histone H3 antibody was used as control for the nucleosomal fraction. *kyp* nucleosomal fraction (time point 0) was used as H3 positive control. **D.** Histogram representing the DDB2 contents at nucleosomes sites (normalized to H3 and time point 0) in WT, j*mj27*, *kyp* and *jmj27kyp* plants prior (0), 30 min and 2h upon UV-C exposure.

Given that DDB2 and JMJ27 load on chromatin 30 min upon UV-C exposure we thus investigated if DDB2 could associate with JMJ27. We generated, by crossing, Arabidopsis plants expressing a functional DDB2-FLAG version (Schalk *et al*., 2017) with either JMJ27-GFP or mJMJ27-GFP (Wang *et al*., 2021a). JMJ27-GFP effectively coimmunoprecipitated with DDB2-FLAG in plant whole-cell extracts prepared before and 30 min upon UV-C exposure whereas mJMJ27-GFP coimmunoprecipitated less (Fig. 6B). These data suggest that DDB2 and JMJ27 form a complex and that the catalytic activity of JMJ27 is important to stabilize this complex upon irradiation.

In human, DDB2 dynamics differs between internucleosomal and core nucleosomal sites (Fei *et al*., 2011). Given that H3K9me2 influences chromatin compaction we analyzed DDB2 kinetics in nucleosomal/internucleosomal fractions prepared from irradiated plants with altered H3K9me2 levels. In WT plants, we found that UV-C irradiation triggers the loading of DDB2 on nucleosomal and on internucleosomal sites at 30 min (Fig. 6C, D). While DDB2 nucleosomal content starts to decrease at 2h, it still increases on internucleosomal sites (Fig. 6C). Hence, 2 pools of DDB2 exist, with different kinetics. Interestingly, *jmj27* plants do not exhibit enrichment of DDB2 in the nucleosomal fraction (Fig. 6C, D). Conversely, in *kyp* and *jmj27 kyp* plants, higher levels of DDB2 are detected along the time course compared to WT and *jmj27* plants (Fig. 6C, D). These results suggest that the H3K9me2 levels regulates DDB2 loading on nucleosomal and internucleosomal sites.

The release of DDB2 from chromatin is crucial for efficient GGR (Molinier *et al*., 2008). Thus, we investigated whether alterations of H3K9me2 contents also interfere with this process. In the WT nucleosomal fraction, DDB2 contents remain stable up to 6h, whilst in *jmj27* plants a decay starts around 2h upon irradiation (Fig. S8A, B). Conversely, in *kyp* plants, increasing amounts of DDB2 are detected up to 6h, suggesting a continuous loading of DDB2 on nucleosomes (Fig. S8A, B). On internucleosomal regions decrease of DDB2 contents starts at 4h in all tested plants (Fig. S8A) highlighting that alteration of H3K9me2 contents does not disturb the kinetics of DDB2 release from internucleosomes.

Thus, these results show that H3K9me2 fine tuning influences DDB2 dynamics at (inter)nucleosomal sites and likely GGR efficiency.

## DISCUSSION

Deciphering the molecular mechanisms regulating accessibility of DNA repair factors to lesions in different chromatin contexts is critical to understand how maintenance of genome integrity occurs along chromosomes. In this study, we combined different approaches to uncover the existence of an interplay between H3K9me2 homeostasis and photodamage repair. We found an important reshaping of chromocenter-like H3K9me2 structures upon UV-C irradiation. Further characterization reveals a disturbance of H3K9me2 reshaping in DDB2 and JMJ27 deficient plants, highlighting that crosstalk between the GGR pathway and the factors regulating H3K9me2 homeostasis might exist. Our genetic approach confirms the existence of such interplay. UV-C-induced loss of H3K9me2 occupancy at chromocenters reflects heterochromatin decompaction. Indeed, the dense constitutive heterochromatin structure hinders an efficient DNA repair process and thus requires decompaction (Schieferstein and Thomä, 1998; Liu, 2015). Alteration of chromatin structure occurs during several DNA repair processes and often results from a change in nucleosome density (Waters *et al*., 2015; Tripuraneni *et al*., 2021; Chakraborty *et al*., 2021). Few emerging models proposed that different chromatin remodeling events facilitate DNA repair, including nucleosome PTM erasure (Palomera-Sanchez *et al*., 2010; Jeon *et al*., 2020), nucleosome sliding and eviction (Dinant *et al*., 2012; Matsumoto *et al*., 2019; Nodelman and Bowman, 2021; Chakraborty *et al*., 2021). Importantly, our results show that constitutive heterochromatin dynamics predominantly depends on the modulation of H3K9me2 content and not of H3K27me1. Although this observation does not exclude the contribution of other mechanisms (*i.e.*, histone eviction), it reflects that histone mark can be specifically/actively removed in an UV-dependent manner. Human cells display an important relaxation of heterochromatin upon UV exposure although no significant decrease of H3K9me3 has been measured (Fortuny *et al*., 2021). Interestingly, in Drosophila, it was shown that the amount of H3K9me3 decreased upon UV exposure, and relies on the histone demethylase KDM4B (Palomera-Sanchez *et al*., 2010). Although it was elusive how the decrease of H3K9me3 is coupled to NER (Palomera-Sanchez *et al*., 2010), our study provides this potential connection. Indeed, we propose a model in which both DDB2 and JMJ27 cooperate as a molecular complex to trigger active H3K9me2 demethylation and chromatin relaxation. This complex would allow DDB2 loading/stabilization at DNA-damaged sites within constitutive heterochromatin. It can be expected that H3K9me2 demethylation is among the first step of chromatin remodeling. In mammals, stabilized DDB2 at damaged sites allows recruiting chromatin remodelers (Pines *et al*., 2012; Wang *et al*., 2021b). Hence, the chromatin relaxation observed upon UV-C exposure may reflect the chain of several mechanisms. Indeed, nucleosome sliding and histone eviction enable the recruitment of the full NER machinery (Zhao *et al*., 2009; Jiang *et al*., 2010; Luijsterburg *et al*., 2012; Pines *et al*., 2012). Interestingly, Quiroz et al. (2024) demonstrated that the mismatch repair factor MUTS HOMOLOG 6 is recruited at H3K4me1 highlighting that histones PTMs play important roles in different DNA repair pathways.

We identified that both KYP and JMJ27 acts in 6-4 PPs repair and in the GGR pathway. These results are in agreement with a recent study showing that GGR is the main pathway for 6-4 PPs excision (Kaya *et al*., 2024). In addition, it highlights that the different types of photolesions are removed by different pathways. This might be due to their formation in genomic regions exhibiting particular epigenetic features that influence the choice and/or the kinetics of the repair pathway (Johann to Berens *et al*., 2022). Indeed, like in human (Fei *et al*., 2011), we uncovered that arabidopsis DDB2 dynamics differs between internucleosomal and core nucleosomal sites, with a crucial influence of H3K9me2 contents. Thus, the control of H3K9me2 homeostasis determines the spatio-temporal dynamics of DDB2 on chromatin.

Analysis of H3K9me2 landscape allowed characterizing that JMJ27 plays a predominant role in the loss of this histone PTM at TEs conversely to IBM1 acting in PCG (Saze *et al*., 2008; Miura *et al*., 2009; Chan and Zimmerli, 2019). To date, JMJ27 was mainly associated with the regulation of gene expression (Dutta *et al*., 2017; Wang *et al*., 2021a). Hence, it cannot be excluded that IBM1 and JMJ27 cooperatively modulate H3K9me2 landscape at PCG and chromocenters, respectively, to fine tune H3K9me2 genome wide under particular growth conditions or during plant development (*i.e.,* meiosis; He *et al*., 2002).

Altogether, these results strengthen the emerging notion in plants that genome/epigenome stability/flexibility are intimately coordinated.

## METHODS

### Plant materials and growth conditions

*Arabidopsis thaliana* plants used in this study are of Col-0 ecotype. Mutant plants are described in Supplemental Table 1 and primers used for genotyping are listed in Supplemental Table 2. Plants were cultivated *in vitro* on solid GM medium [MS salts (Duchefa), 1% sucrose, 0.8% Agar-agar ultrapure (Merck), pH 5.8] and grown in a culture chamber under a 16 h light (light intensity ∼150 μmol m^−2^ s^−1^; 21°C) and 8 h dark (19°C) photoperiod.

### UV-C irradiation

21-day-old *in vitro* grown arabidopsis plants were exposed to 3,000 J/m^2^ of UV-C using the Stratalinker 2400 (Stratagene). Plant material was harvested prior irradiation for control (0) and at different time points upon irradiation (30 min, 2h, 4h, 6h or 24h) for further processing.

### UV-C sensitivity assay

UV-C (λ = 254 nm) sensitivity was evaluated on 6-day-old *in vitro*-germinated arabidopsis plants. Plants were transferred to square plates containing GM medium and grown vertically. Root length was measured 24h upon UV-C exposure (300, 600 and 900 J/m^2^) using the Stratalinker 2400 (Stratagene). The relative root growth was calculated as followed: (root length treated/root length untreated) × 100 (± SD). Eight plants per replicate were used. Experiments were performed in triplicates.

### Photodamage removal assay

Twenty-one-day-old *in vitro* arabidopsis grown seedlings (n=40 per genotype) were irradiated with UV-C (3,000 J/m^2^). Half of the samples was harvested immediately after irradiation (time 0) and the other half was kept under dark condition for 1h to prevent photolyases to revert photoproducts. Genomic DNA was extracted using plant DNA extraction kit (Macherey-Nagel). DNA samples were processed as described in Schalk *et al*. (2017). Repair efficiency was determined by the quantification of the remaining CPDs or 6-4 PPs amounts after 1h, relative to the photodamage content at time 0.

### Immunolocalization of photolesions

Leaves 3 and 4 of 21-days old *in vitro* grown arabidopsis plants were harvested prior irradiation for control (0) and 30 min upon irradiation. Samples were incubated 4 times at least 5 min in fixative cold (4°C) solution (3:1 ethanol/acetic acid; vol/vol). Leaves nuclei were extracted by chopping fixed tissue in LB-01 Buffer (15 mM Tris-HCl pH 7.5, 2 mM EDTA, 0.5 mM spermine, 80 mM KCl, 29 mM NaCl, 0.1% Triton X-100) with a razor blade. The nuclei containing solution was filtered through 20 µm nylon mesh and centrifugated 1 min (1,000g). Supernatant was spread on poly-lysine slides (Thermofisher: 631-1349) and post fixation was performed using a 1:1 acetone / methanol (vol/vol) solution for 2 min. Slides were washed with Phosphate Buffer Saline x1 and incubated for 1h at room temperature in permeabilization buffer (8% BSA, 0.01% Triton-X in Phosphate Buffer Saline x1). Slides were incubated over night at 4°C with anti-CPDs or anti-6-4 PPs primary antibodies (CosmoBio: CAC-NM-DND-001; CAC-NM-DND-002) diluted in 1% BSA, Phosphate Buffer Saline x1 buffer.

### Immunolocalization of histone marks

Leaves 3 and 4 of 21-days old *in vitro* grown arabidopsis plants were harvested prior irradiation for control (0) and 2h, 24h upon irradiation. Samples were fixed at 4°C in paraformaldehyde solution (4% in PBS). Leaves nuclei were extracted by chopping fixed tissue in LB-01 Buffer (15 mM Tris-HCl pH 7.5, 2 mM EDTA, 0.5 mM spermine, 80 mM KCl, 29 mM NaCl, 0.1% Triton X-100) with a razor blade. The nuclei containing solution was filtered through 20 µm nylon mesh and centrifugated 1 min (1,000g). Supernatant was spread on poly-lysine slides (Thermofisher: 631-1349) and post fixation was performed using paraformaldehyde (3.2% in PBS). Slides were washed with Phosphate Buffer Saline x1 and incubated for 1h at room temperature in permeabilization buffer (8% BSA, 0.01% Triton-X in Phosphate Buffer Saline x1). After the permeabilization step, slides were incubated over night at 4°C with primary-antibody (H3K9me2: Diagenode: C15200154; H3K27me1: Diagenode: C15410045-50) diluted in 1% BSA, Phosphate Buffer Saline x1 buffer.

### Secondary antibodies for immunolocalization

Upon incubation slides were washed at least 3 times with PBS. Secondary antibody coupled to Alexa fluor 488 (ThermoFisher: A-11001) or Alexa fluor 568 (ThermoFisher: A-11004), diluted in 1% BSA, PBS, was added and incubated for 90 min at room temperature. Finally, slides were washed 3 times with PBS and 15 μl of Fluoromount-G (Southern Biotechnology: 0100–01) with 2 μg/ml DAPI were added as mounting solution for the coverslip.

### Microscopy Image acquisition, segmentation and measurements

Image acquisition was entirely performed on a Zeiss LSM 780 confocal microscope using a 64X oil immersion objective. A 405 nm, 488 nm and 568 laser excitation wavelengths were used for DAPI, Alexa Fluor 488/GFP and Alexa Fluor 526, respectively. Emission DAPI was measured considering wavelengths in the range 410-585. Alexa Fluor 488/GFP emission was measured considering wavelengths in the range 493-630 nm. Alexa Fluor 568 emission was measured considering wavelengths in the range 590-645 nm. The same acquisition gain settings were used for all slides of a same experiment. Each image acquisition consists in a Z-stack capture with a 0.64 μm slice distance. Regions of interest were segmented by the deep learning script Nucl.Eye.D (https://zenodo.org/records/7075507) allowing the quantification of the following nuclear features:

- PTM Occupancy: percentage of surface occupied by all bright immunolabelled chromocenters-like structure in the corresponding nucleus.
- Relative PTM intensity: ratio of mean chromocenter-like structure intensity / Mean nucleus intensity
- PTM Content (Occupancy * Relative Intensity)

### Chromatin preparation and Mnase treatment

Leaves 3 and 4 of 21-days old *in vitro* grown arabidopsis plants were harvested prior irradiation for control (0), 30 min, 2h, 4h and 6h upon irradiation. Fractions of soluble (S1)/insoluble (P1) proteins were extracted from 150 mg of plant material using Nonidet P-40 lysis buffer (25 mM Tris-HCl, pH 8.0, 0.3 M NaCl, 1 mM EDTA, 10% [v/v] glycerol, Nonidet P-40 1% [v/v], 0.2 mM phenylmethylsulfonyl fluoride, and EDTA-free Protease Inhibitor Cocktail [1 tablet/50 mL]). After grinding, powder was resuspended in 2.5 ml of Nonidet P-40 lysis buffer and incubated for 30 min on a rotating wheel at 4°C (8 rpm) and the solution was filtered through Miracloth. Free chromatin-unbound proteins (S1 fraction) were recovered from the insoluble chromatin-enriched fraction (P1) after centrifugation (13,000g, 10 min, 4°C). S1 and P1 fractions were loaded in denaturing buffer for SDS-PAGE separation and immunoblotted with the indicated antibodies (anti-DDB2: Molinier *et al*., 2008; anti-GFP: Takara-632593; anti-histone H3: Agrisera-AS10710; Anti-UGPase: Agrisera-AS142813). The pellet containing insoluble and chromatin-bound proteins (P1) was washed twice with 0.5 ml ice-cold CS buffer (20 mM Tris-HCl, pH 7.5, 100 mM KCl, 2 mM MgCl2, 1 mM CaCl2, 0.3 M sucrose, 0.1% (v/v) Triton X-100) for Mnase treatment. This P1 fraction was resuspended in 40 µl of CS buffer. Mnase treatment was performed by adding 5 µl of 10× reaction buffer [500 mM Tris-HCl (pH 7.9), 50 mM CaCl2], 1 µl of bovine serum albumin (BSA; 1 mg/ml) and MNase (4 U/µl in a volume of 50 µl) and incubated at 37 °C for 15 min. MNase digestion was stopped by the addition of EGTA (5 mM) and the internucleosomal fraction proteins (S2) was separated from insoluble core nucleosome fraction (P2) by centrifugation at 15, 000 g (10 min, 4 °C). P2 fraction was resuspended in 50 µl of CS buffer. S2 and P2 fractions were loaded in denaturing buffer for SDS-PAGE separation and immunoblotted with the indicated antibodies (anti-DDB2: Molinier *et al*., 2008; anti-histone H3: Agrisera-AS10710, anti-H3K9me2: Abcam-ab176882).

### Co-immunoprecipitation

21-days old *in vitro* grown arabidopsis plants were harvested prior irradiation for control (0), and 30 min upon UV-C irradiation. Proteins were extracted from 200 mg of plant material using 3 ml of IP buffer (50 mM Tris-HCl at pH 7.5, 150 mM NaCl, 5 mM MgCl2, 0.1% v/v NP40, 10% glycerol, EDTA-free Protease Inhibitor Cocktail [1 tablet/50 ml]) and incubated for 30 min on a rotating wheel at 4°C (8 rpm). The solution was Miracloth-filtered and immunoprecipitation was performed using anti-FLAG gel affinity (Sigma-Aldrich) over night on a rotating wheel at 4°C (8 rpm). The precipitate was washed 4 times in IP buffer, resuspended in 50 μl of denaturing buffer and heated for 5 min at 95°C prior to immunoblotting. The DDB2-FLAG fusion protein was detected using the anti-FLAG HRP (Sigma-Aldrich: A8592) at a 1:5,000 (v: v) dilution in PBST (1× PBS, nonfat dry milk [5%, w/v], and Tween 20 [0.1%, v/v; Sigma-Aldrich]). The JMJ27-GFP fusion protein was detected using the anti-GFP (Takara: 632593) 1:2,000 (v: v) dilution. The anti-Rubisco large subunit antibody (Agrisera: AS03037) was used as loading control.

### RNA extraction and RT-qPCR

Total RNAs were extracted from untreated (0) and UV-C treated WT and *jmj27* plants (time points 30 min, 1h and 2h) using Tri-Reagent (Sigma). Reverse transcription reaction (RT) was performed on 5 µg of total RNA using the SuperScript IV (Thermo Fischer) with a mixture of random hexamers-oligo d(T) primers. 100 ng of the RT reaction was used for quantitative PCR (qPCR). qPCR was performed with technical triplicates, using a Light Cycler 480 and Light Cycler 480 SYBR green I Master mix (Roche) following manufacturer’s instructions. All primers are listed in Supplemental Table 2. Experiment was duplicated using independent biological replicates.

### Chromatin immunoprecipitation

21-days old *in vitro* grown arabidopsis plants (WT and *jmj27*) were harvested prior irradiation for control (0), 30 min and 2h upon UV-C irradiation. Two independent biological replicates have been performed with 70 plants per replicate and time point. ChIP was performed using the Universal Plant Chip-seq Kit (Diagenode: C01010152) according manufacturers’ instructions and H3K9me2 (Diagenode: C15200154), H3 (Abcam: ab10799) antibodies. Libraries were prepared with the MicroPlex Library Preparation Kit v3 (Diagenode: C05010001) and the 24 Dual indexes for MicroPlex Kit v3 (Diagenode-: C05010003). Illumina single end sequencing was subsequently performed (1×100) on a NextSeq 2000 System (Illumina).

### ChIP-seq analysis

ChIP-seq analysis has been performed using the community curated nf-core ChIP-seq pipeline (version 1.2.2; Ewels *et al*., 2020) with default parameters. Reads have been mapped on the reference genome (TAIR10) and MACS2 (Zhang *et al*., 2008) was used to call H3K9me2 broad peaks using H3 as a reference, and an effective genome size of 119,481,543 bp. Replicates were handled using the stringent irreproducibility discovery rate (IDR) framework (Li *et al*., 2011) and only peaks containing an IDR value superior or equal to a scaled IDR value of 540 were considered real, corresponding to an IDR of 0.05. Statistics of ChIP-sequencing are shown in Supplemental Table 3. Venn diagrams have been produced using the R package “VennDiagram”. JMJ-27-dependent loci have been identified by the comparison of the set of genomic regions loosing H3K9me2 signal(s) at time point 30 min and/or 2h in WT plants with the genomic regions in which the H3K9me2 signal is maintained at time point 30 min and/or 2h in *jmj27* plants. A minimal overlap of 200 bp between regions was applied.

### Statistics

Mann-Whitney U or Wilcoxon Matched-Pairs Signed-Ranks tests were used as non-parametric statistical hypothesis tests (http://astatsa.com/WilcoxonTest/). Chi 2 test was used to determine significant difference between categories distribution (https://goodcalculators.com/chi-square-calculator/). t-test was used as parametric statistical hypothesis test.

## Acknowledgments

We are grateful to Prof. Jin for providing the JMJ27-GFP and mJMJ27-GFP seeds. We acknowledge the IBMP Cell Imaging Facility, member of the national infrastructure France-BioImaging supported by the French National Research Agency (ANR-10-INBS-04). We thank Anne Molitor, Antoine Hanauer and Raphaël Carapito (Institut National de la Santé et de la Recherche Médicale, GENOMAX platform, UMRS1109) for Next-Generation sequencing experiments. This research was funded by a grant from the French National Research Agency (ANR-20-CE20-002) and supported by the EPIPLANT Groupement de Recherche (CNRS, France).

## Data availability

All sequencing data used in this study have been deposited at NCBI under the XXXX and will publicly available as of the date of publication.

## Code availability

The original code for Nucl.Eye.D is available at https://zenodo.org/records/7075507. Any additional information is available from the corresponding author upon request.

---

**Supplemental Table 1:** *Arabidopsis thaliana* mutant lines used in the study.

| **Mutant** | **Reference** |
| --- | --- |
| *phr1* | WiscDsLox466C12 |
| *uvr3* | WiscDsLox334H05 |
| *ddb2-3* | WiscDsLoxHs19505H |
| *jmj27* | SAIL400B08 |
| *ibm1-6* | SALK006042 |
| *kyp* | SALK069326 |
| *kyp suvh5/6* | SALK041474 GK-263C05 SAIL1244F04 |

**Supplemental Table 2:**
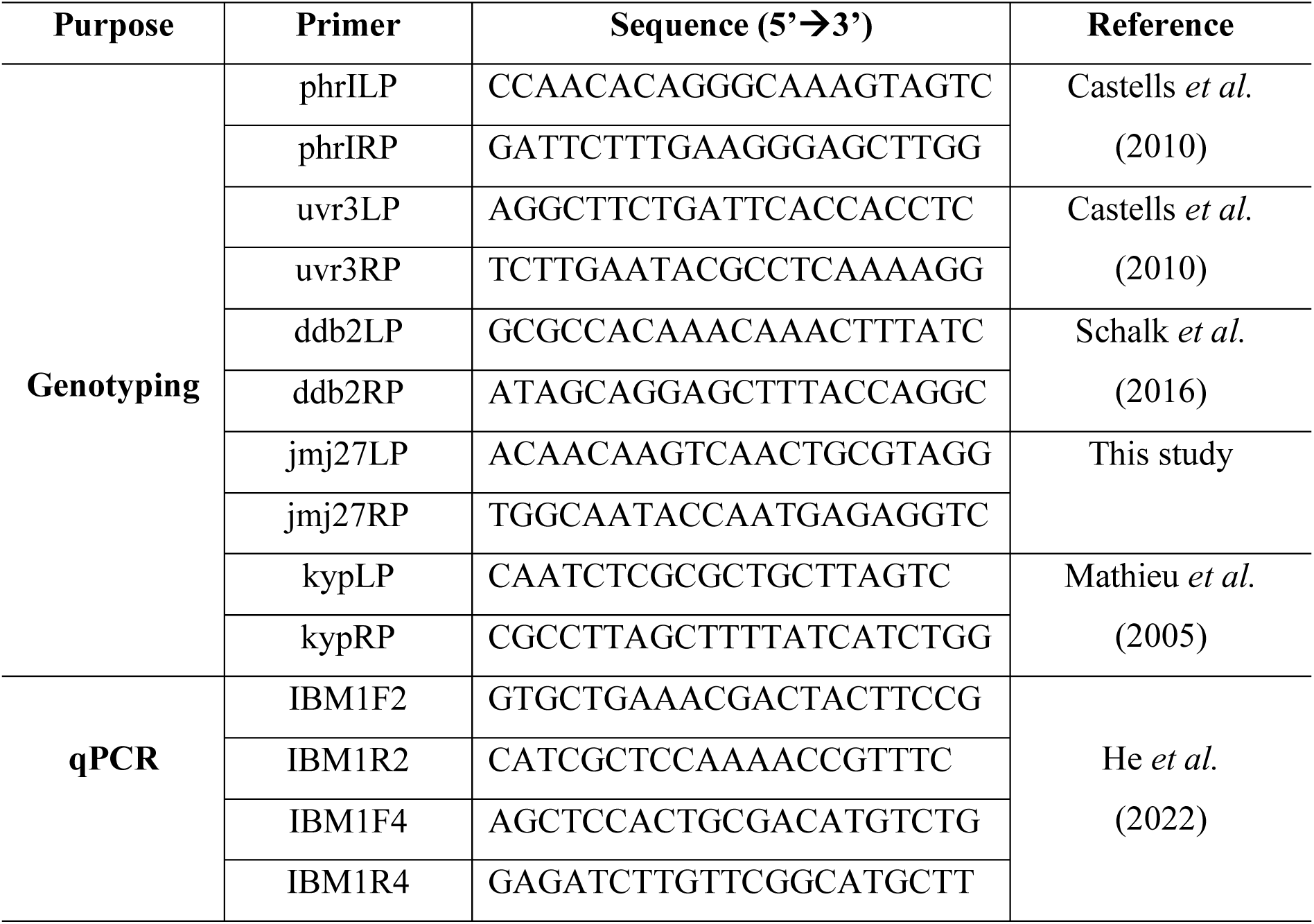
Primers used in the study.

**Supplemental Table 3:** Statistics of the ChIP-sequencing experiments.

| Genotype | Time point | Antibody | Biological replicate | Total number of reads | Mapped reads |
| --- | --- | --- | --- | --- | --- |
| WT | 0 | H3 | 1 | 16295137 | 8978322 |
| WT | 0 | H3 | 2 | 15414793 | 7388192 |
| WT | 30 min | H3 | 1 | 17825994 | 6234466 |
| WT | 30 min | H3 | 2 | 17879995 | 10784977 |
| WT | 2h | H3 | 1 | 17025272 | 7160201 |
| WT | 2h | H3 | 2 | 16906111 | 10011941 |
| WT | 0 | H3K9me2 | 1 | 21942394 | 4605327 |
| WT | 0 | H3K9me2 | 2 | 22695011 | 2702865 |
| WT | 30 min | H3K9me2 | 1 | 22206619 | 6572873 |
| WT | 30 min | H3K9me2 | 2 | 20649852 | 9132343 |
| WT | 2h | H3K9me2 | 1 | 22468208 | 7165852 |
| WT | 2h | H3K9me2 | 2 | 23158430 | 3143648 |
| *jmj27* | 0 | H3 | 1 | 15262557 | 9289119 |
| *jmj27* | 0 | H3 | 2 | 13451293 | 7412342 |
| *jmj27* | 30 min | H3 | 1 | 17772710 | 6961404 |
| *jmj27* | 30 min | H3 | 2 | 16538338 | 6110753 |
| *jmj27* | 2h | H3 | 1 | 16114361 | 9001334 |
| *jmj27* | 2h | H3 | 2 | 15665448 | 6110753 |
| *jmj27* | 0 | H3K9me2 | 1 | 19928173 | 9500876 |
| *jmj27* | 0 | H3K9me2 | 2 | 18763748 | 8174707 |
| *jmj27* | 30 min | H3K9me2 | 1 | 20073922 | 6703878 |
| *jmj27* | 30 min | H3K9me2 | 2 | 22248110 | 4442291 |
| *jmj27* | 2h | H3K9me2 | 1 | 20194725 | 6269076 |
| *jmj27* | 2h | H3K9me2 | 2 | 18695805 | 6713164 |

**Supplemental figure 1:**
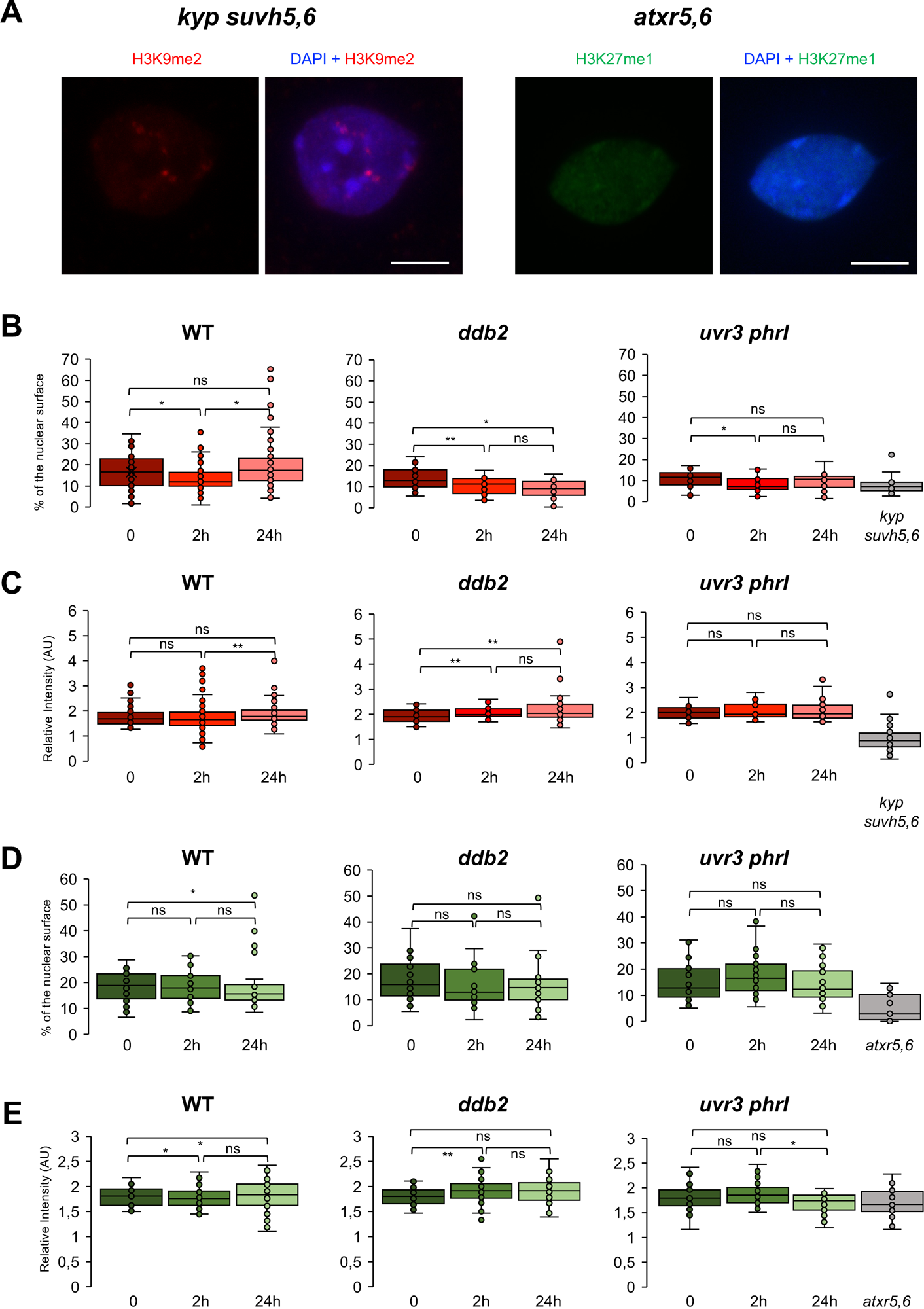
**A.** Microscopy images of (top) DAPI and H3K9me2 immuno-stained arabidopsis nuclei isolated from *kypsuvh5/6* and of (bottom) DAPI and H3K27me1 immuno-stained arabidopsis nuclei isolated from *atxr5/6* leaves in control condition. **B.** Box plots of H3K9me2 occupancy (percent of the nuclear surface occupied by chromocenter like structure). **C.** Same as **B.** for H3K27me1 occupancy. **D.** Box plots of H3K9me2 Relative Intensity (ratio of mean chromocenter structure intensity / Mean nuclei intensity). **E.** same as **D.** for H3K27me1 Relative Intensity. A population of at least 25 nuclei per time point was used. Each dot represents the measure for 1 nucleus. Mann Whitney Wilcoxon test \**p* < 0.01; \*\**p*< 0.05 ns: nonsignificant.

**Supplemental figure 2:**
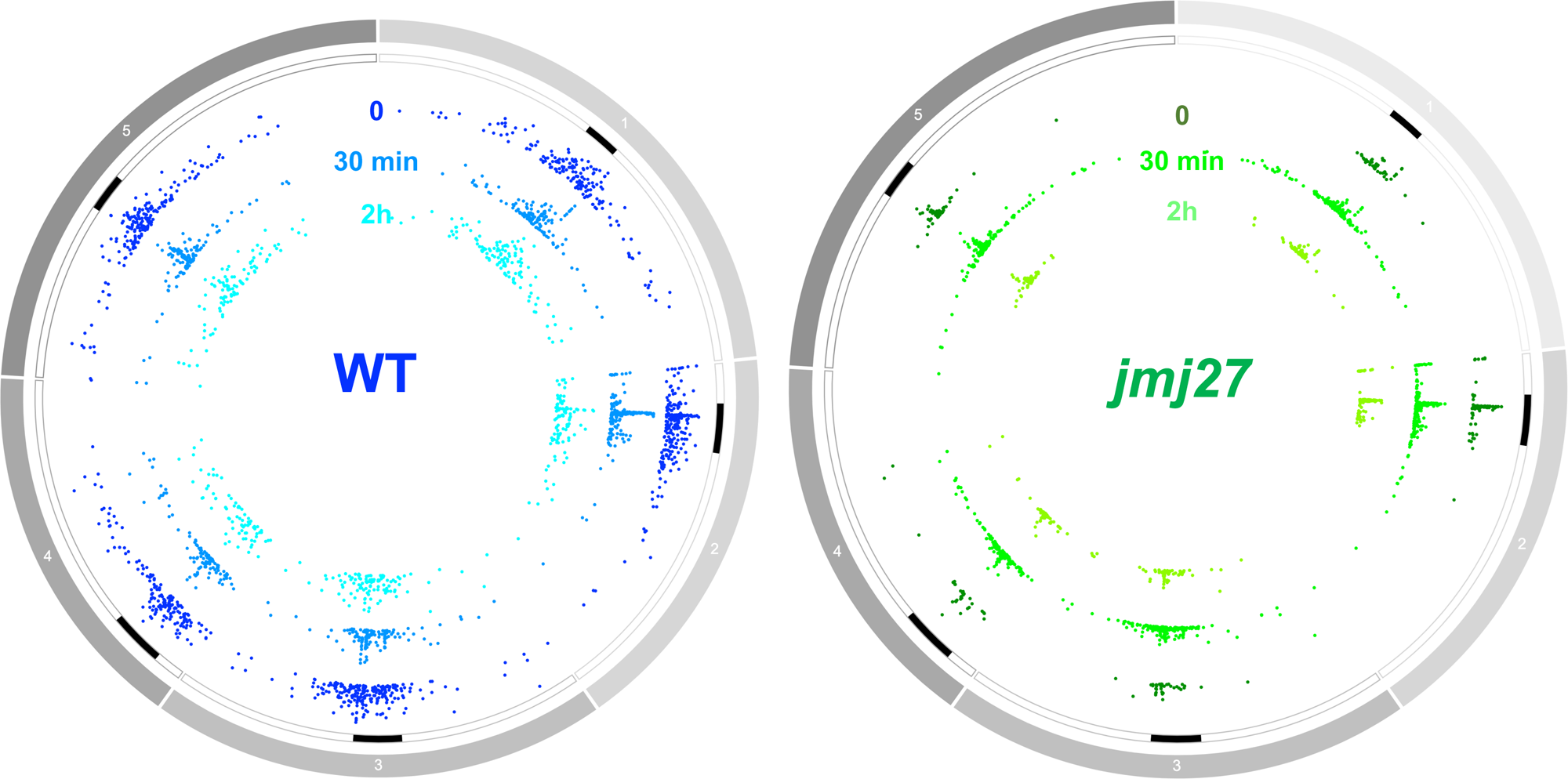
Genome wide distribution of H3K9me2-enriched regions. Circos representation of H3K9me2 landscape in WT (left) and in *jmj27* (right) plants at different time points upon UV-C exposure. Each chromocenter is shown by a black rectangle.

**Supplemental figure 3:**
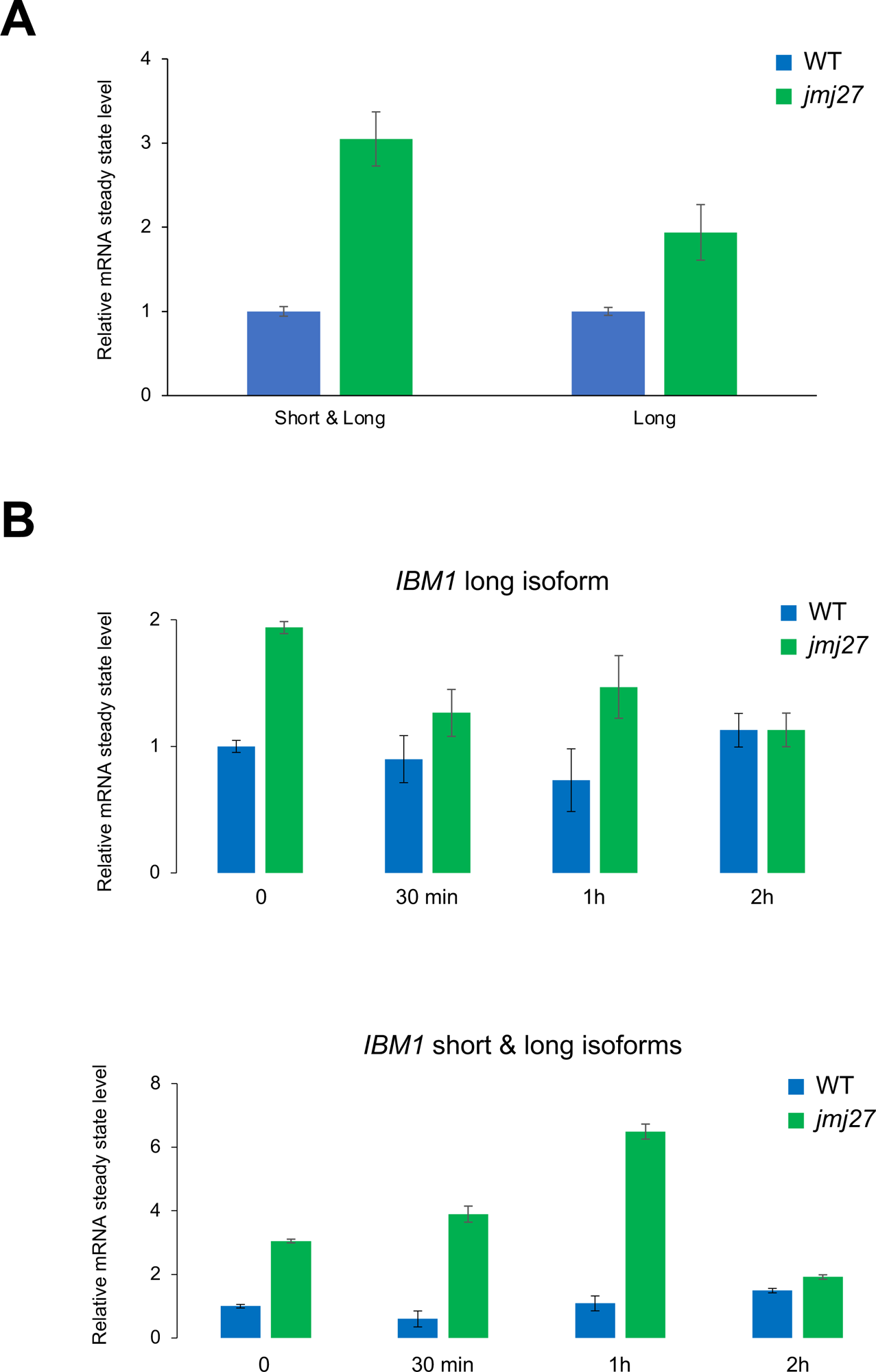
*IBM1* mRNA levels. Relative mRNA steady state level (±SD) of long and short *IBM1* isoforms **(A)** in *jmj27* plants relative to WT plants and **(B)** during a time course following UV-C exposure in WT and *jmj27* plants (relative to WT time point 0).

**Supplemental figure 4:**
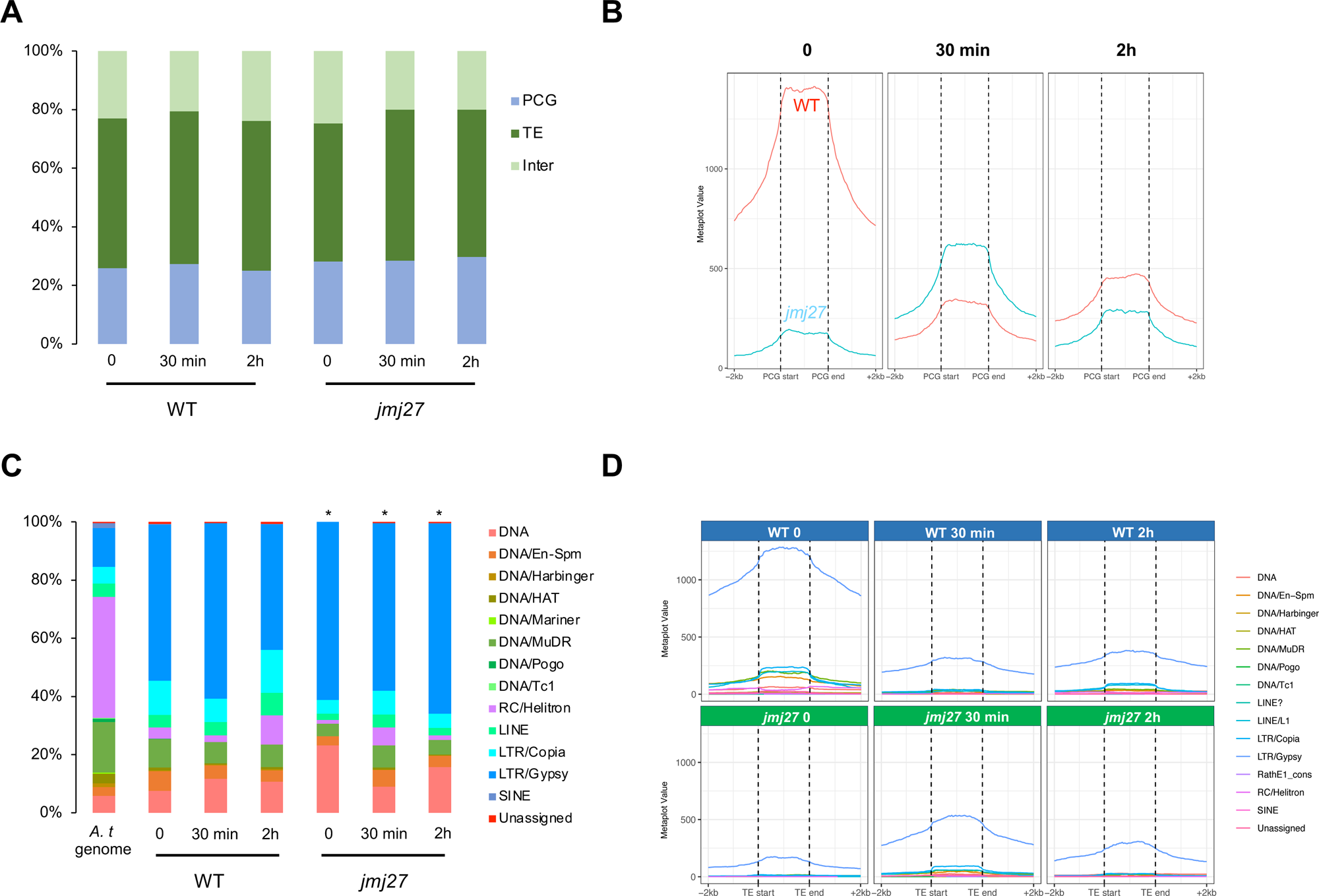
Genetic elements enriched in H3K9me2. **A.** Histogram showing the distribution of the genetic elements (Protein Coding Genes: PCG; Transposable Elements: TE and Intergenic regions: Inter) enriched in H3K9me2 in WT and *jmj27* plants, prior (0) and upon UV-C exposure (30 min and 2h). **B.** Metaplots showing the average H3K9me2 signal at PCG. **C.** Histogram showing the distribution of the TE superfamilies enriched in H3K9me2 in WT and *jmj27* plants, prior and upon UV-C exposure. *A.t* genome represents the distribution of TE superfamilies in the *Arabidopsis thaliana* genome. * Chi square test < 0.01 compared to the corresponding time point in WT plants. **D.** Same as **B.** for TE superfamilies. TEs and PCGs were aligned at their 5′ and 3′ ends (dashed lines). Average methylation over 100 bp bins 2kb upstream and 2kb downstream from the alignment points is plotted.

**Supplemental figure 5:**
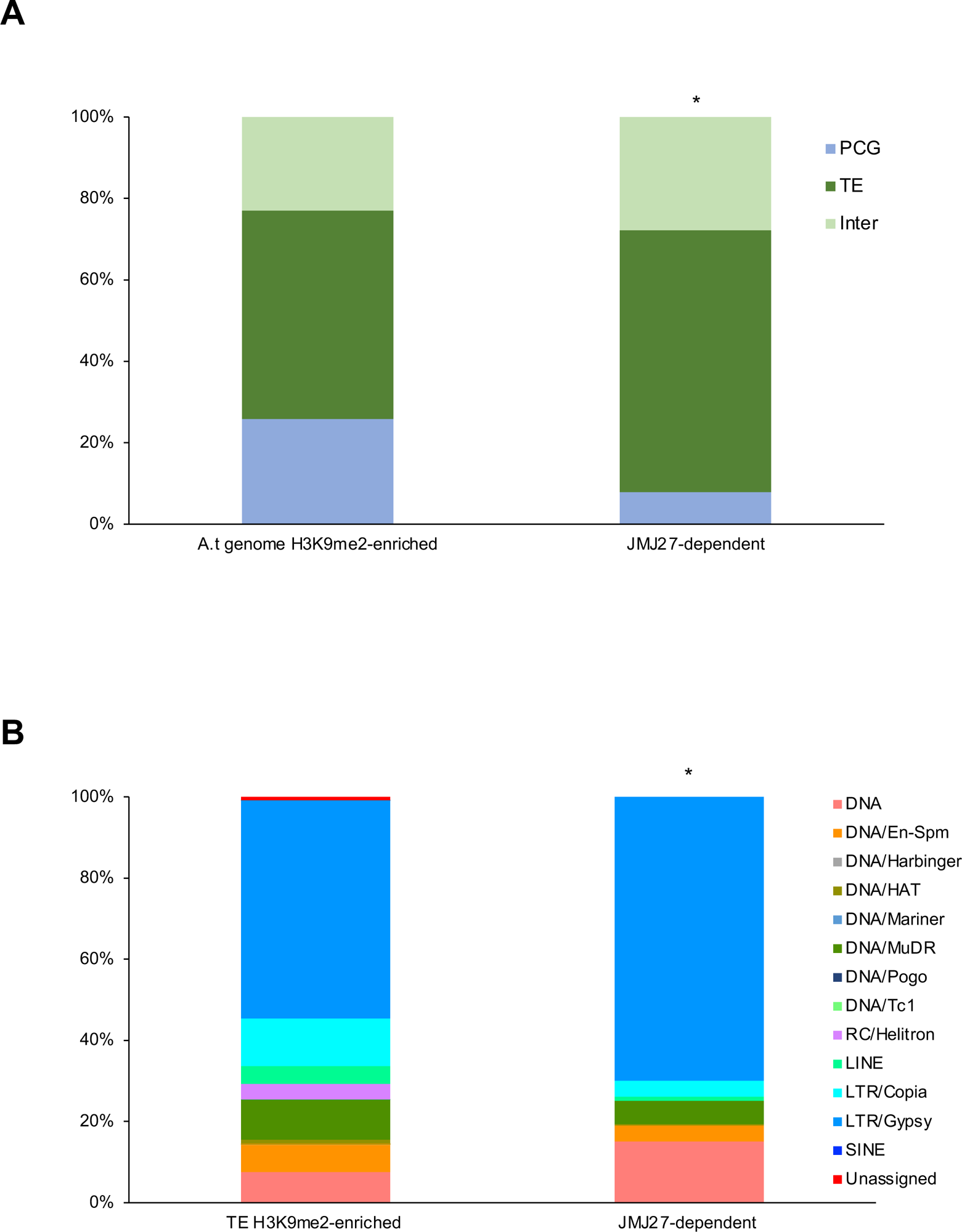
JMJ27-dependent genomic regions. **A.** Histogram showing the distribution of the JMJ27-dependent genetic elements (Protein Coding Genes: PCG; Transposable Elements: TE and Intergenic regions: Inter). A.t genome enriched in H3K9me2 at PCG, TE and Intergenic regions based on our ChIP experiments (WT 0). * Chi square test < 0.01 compared to the *A*. *t*. genome enriched in H3K9me2. **B.** Histogram showing the distribution of the JMJ27-dependent TE superfamilies. * Chi square test < 0.01 compared the H3K9me2-enriched *A*. *t*. genome/TE.

**Supplemental figure 6:**
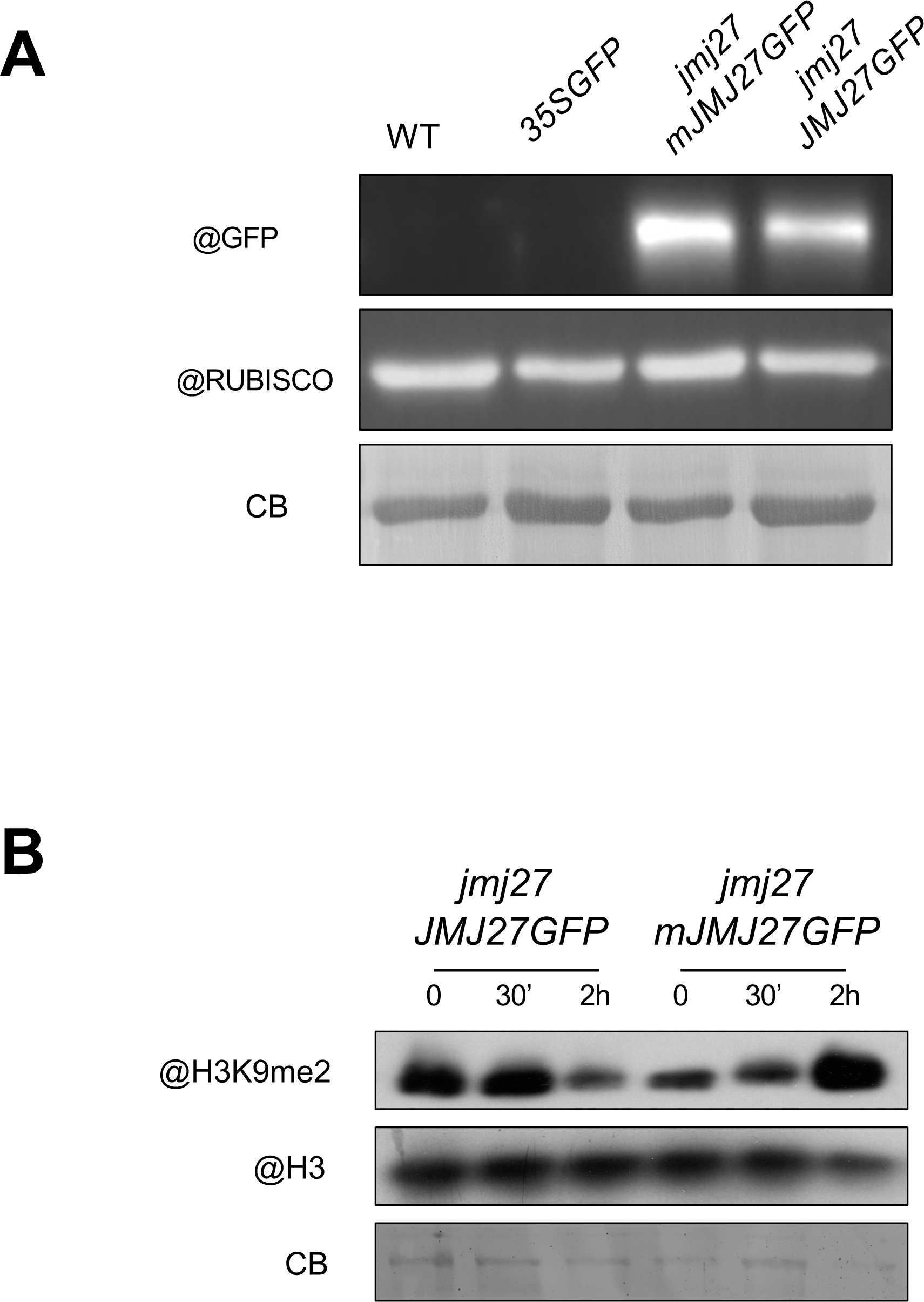
*jmj27*-complemented plants. **A.** Immunoblot analysis of JMJGFP and mJMJGFP protein contents (125kDa) in *jmj27 JMJ27GFP* and *mJMJ27GFP* plants. WT and *35SGFP* plants were used as negative control. Coomassie blue staining (CB) of the blot is shown. **B.** Immunoblot analysis of H3K9me2 contents in the nucleosomal fraction of *jmj27 JMJ27GFP* and *mJMJ27GFP* plants prior (0), 30 min and 2h upon UV-C exposure. Anti-H3 antibody was used as loading control. Coomassie blue staining (CB) of the blot is shown.

**Supplemental figure 7:**
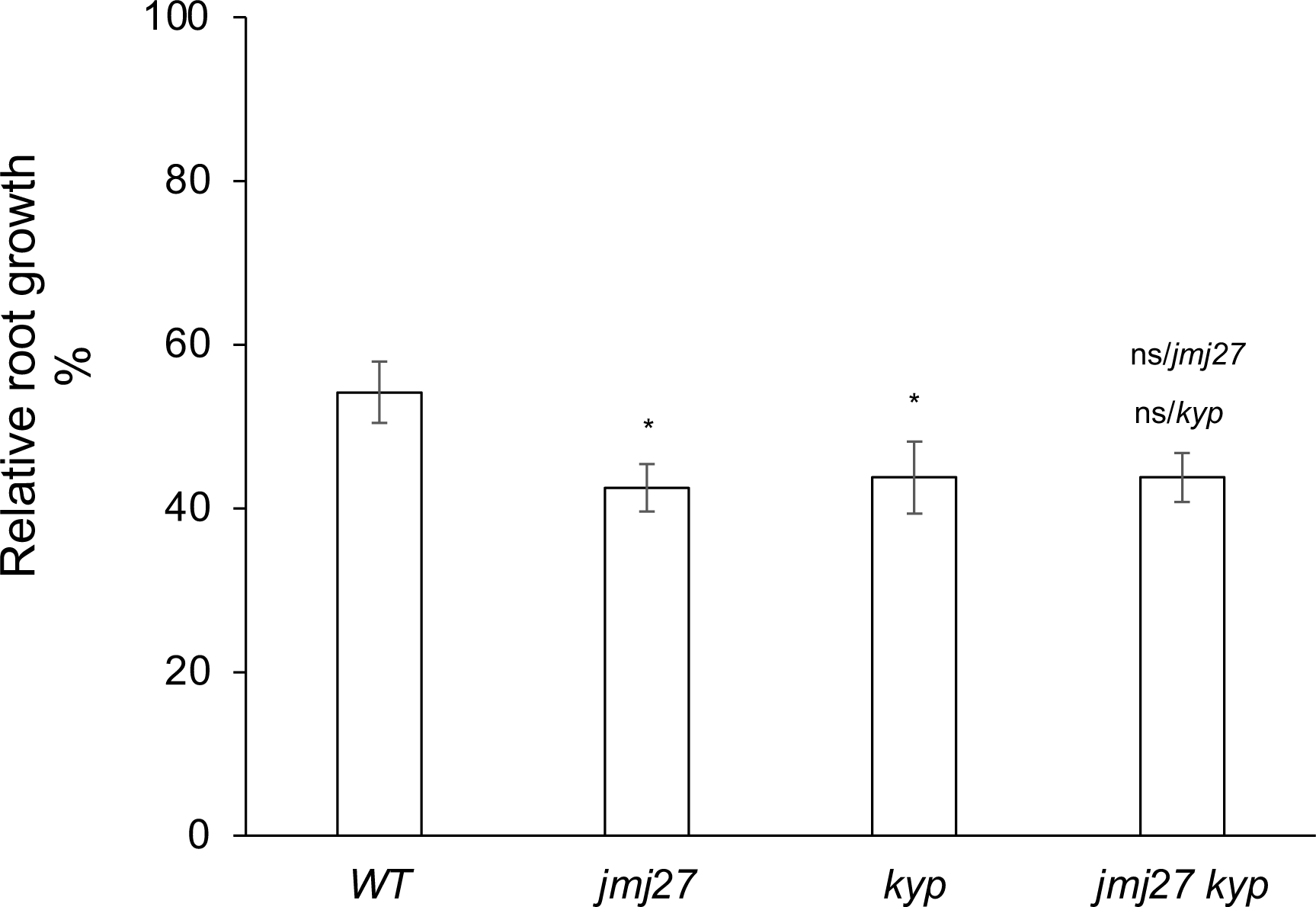
Genetic interactions between *jmj27* and *kyp*. UV-C sensitivity assay of WT, *jmj27, kyp* and *jmj27 kyp* plants using the root growth assay. *t* test \**p* < 0.01; ns, nonsignificant compared with WT plants or with the single mutant plant. Eight plants per replicate were used, and three independent biological replicates were performed.

**Supplemental figure 8:**
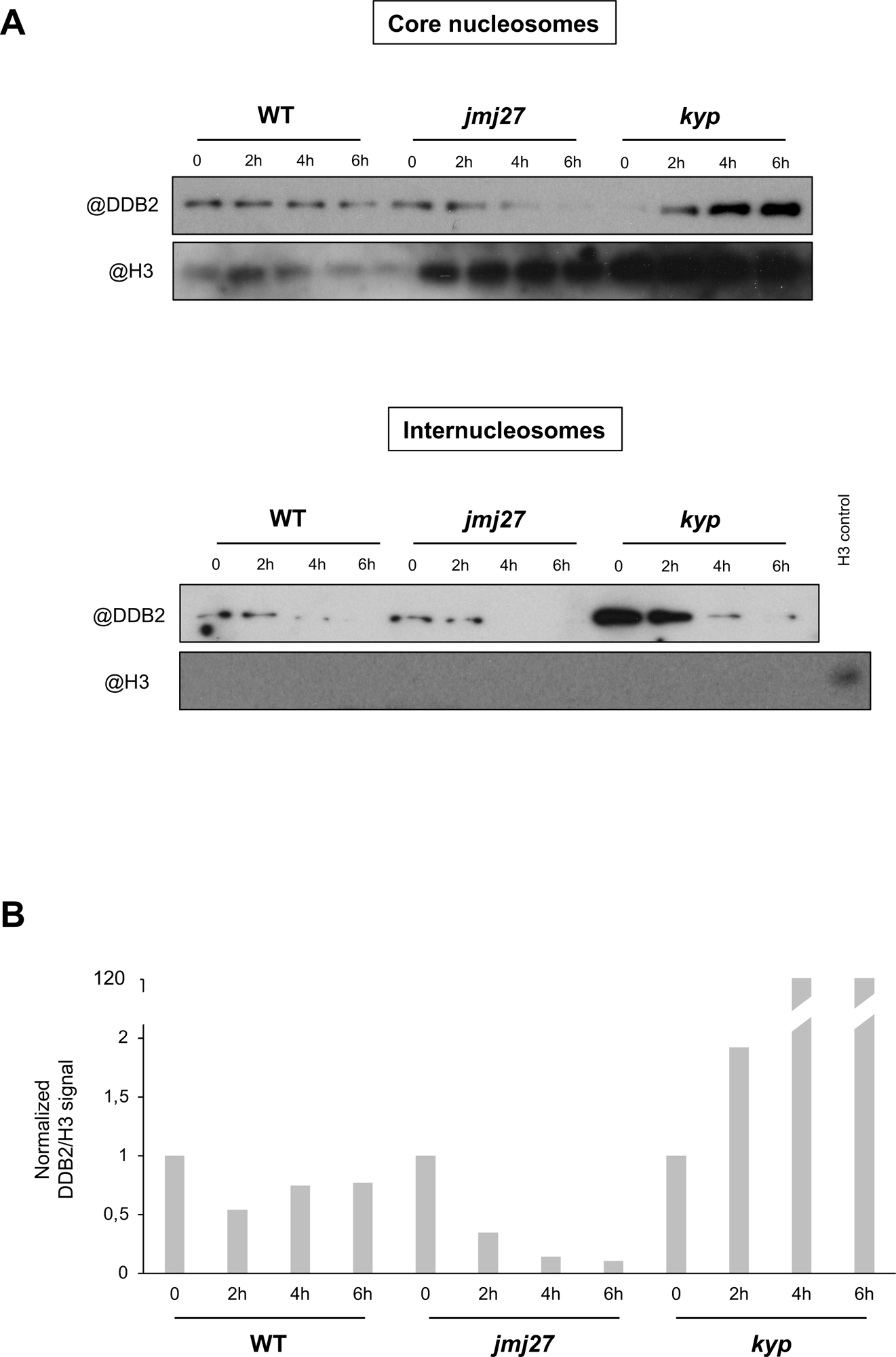
DDB2 contents at internucleosomal and nucleosomal sites. **A.** Immunoblot analysis of DDB2 content at nucleosomal (top panel) and at internucleosomal (bottom panel) sites in WT, j*mj27* and *kyp* plants prior, 2h, 4h and 6h upon UV-C exposure. Anti-histone H3 antibody was used as control for the nucleosomal fraction. *kyp* nucleosomal fraction (time point 0) was used as H3 positive control **B.** Histogram representing the DDB2 contents at nucleosomes sites (normalized to H3 and time point 0) in WT, j*mj27* and *kyp* plants prior (0), 2h, 4h and 6h upon UV-C exposure.

## Notes

### Competing Interest Statement

The authors have declared no competing interest.

